# Cofilin controls actin network identity by sorting actin binding proteins to distinct cytoskeletal structures

**DOI:** 10.64898/2026.06.20.733546

**Authors:** Dominic M. Radcliffe-Hines, Ruben Bonilla Santiago, Alexander J.B. Reading, Brian S. Hercyk, Carys R. Evans, Shane G. McInally

## Abstract

Proper cell physiology requires the co-assembly of multiple actin cytoskeletal networks that are tailored for specific functions. To maintain and promote the different functions of these networks, cells decorate them with distinct types of actin binding proteins (ABPs). While various models have been proposed to explain this selective sorting of ABPs, the role of actin disassembly factors is less well understood. Here, we used inducible CRISPR interference and quantitative live-cell imaging to test how disassembly factors control the ABP composition of different networks. We found that knockdown of cofilin (Cof1), a potent and highly conserved disassembly factor, disrupts the size, organization, and ABP composition of actin networks. Specifically, defects in Cof1-mediated disassembly disrupt intracellular transport due to the assembly of overgrown and disordered branched actin networks that are inappropriately decorated by tropomyosin (Tpm1). Contrary to prevailing models of ABP sorting, these networks are co-decorated by Tpm1 and fimbrin (Sac6), and their assembly is independent of formin activity. Instead, our findings support a model wherein failure to maintain the proper architecture of branched actin networks drives mis-localization of network-specific ABPs. Together, this work demonstrates that actin disassembly factors play a critical role in maintaining cytoskeletal structure and function to regulate ABP sorting across distinct networks.

## Introduction

Eukaryotic cells build multiple, functionally diverse actin cytoskeletal networks at the same time and within the same cytoplasm. These networks are built from a common pool of molecular building blocks, yet have distinct architectures and dynamics that are tailored for their specific cellular functions^1,2^. The assembly of networks with distinct architectures is driven by actin nucleators – the Arp2/3 complex promotes assembly of branched actin networks, while formins and Ena/VASP nucleate linear networks^3^. Further control over the size, shape, and dynamics of these networks is accomplished through the selective association of distinct sets of actin binding proteins (ABPs)^4,5^. Therefore, sorting ABPs to the correct types of actin networks is critical for establishing and maintaining actin network identity and function. However, it remains unclear how cells differentiate between coexisting actin structures to ensure that each is decorated by the correct ABPs.

Current models used to explain this important behavior are based on competitive and cooperative binding of ABPs to either the sides of actin filaments, or to their growing barbed ends^5^. In these models, sorting is driven by complex interactions between different ABPs that result in the segregation of these regulators to distinct networks^4,5^. For example, fimbrin and tropomyosin are thought to directly compete for binding to the sides of actin filaments, driving their exclusive association with branched and linear networks, respectively^6,7^. Capping protein and formins are also thought to compete for binding to barbed ends, leading to the exclusive association of capping protein with Arp2/3-generated branched actin networks^8,9^. In both cases, disrupting the competition between ABPs leads to sorting defects associated with compromised network assembly and function. Specifically, tropomyosin is inappropriately recruited to branched networks in fimbrin and capping protein mutants, leading to defects in endocytosis and intracellular transport, respectively^6,8,9^. In addition to these competition mechanisms, intrinsic differences in the filaments assembled by different nucleators may also contribute to controlling which ABPs associate with different actin networks^10,11^.

Another critical feature of physiological actin networks is the ability to rapidly turnover^3,12^. This allows cells to quickly rearrange and reorganize their actin cytoskeleton in response to stimuli and depends on the activity of disassembly factors that increase the rate of turnover. Beyond enabling network plasticity, robust turnover ensures that cells maintain an available pool of monomers to support continuous filament assembly and preserve the homeostatic balance between branched and linear actin networks^13^. Disassembly factors are thought to orchestrate network turnover by collaborating through distinct mechanisms that promote severing, depolymerization, or debranching of filaments^12,14^. In this way, a single disassembly factor may participate in multiple disassembly pathways, functioning either independently or in collaboration with other factors to drive network turnover.

The turnover rate of actin networks is also shaped by their ABP composition, since ABPs can either prevent or promote the association of additional regulators. For instance, tropomyosin can prevent cofilin-mediated severing of filaments^6,15^, slowing the rate at which they turnover. Coupling these inhibitory interactions with the competitive exclusion of tropomyosin from fimbrin-decorated networks renders some filament populations susceptible to cofilin’s disassembly activity while insulating others. Further, in vitro work has demonstrated that competition between network-specific ABPs, such as cofilin and fimbrin, can synergize rather than inhibit their individual functions^7^. While the interactions between ABPs and disassembly factors are clearly important for proper network assembly and turnover, it remains unknown whether disassembly factors actively control ABP sorting, or whether their activities are simply tuned by the existing ABP composition of a given network.

Budding yeast are a powerful system to dissect the mechanisms that control ABP sorting and define actin network identity. During polarized growth, these cells continuously build two functionally distinct actin networks: Arp2/3-generated cortical patches that perform endocytosis, and formin-generated polarized cables that serve as tracks for myosin-based intracellular transport^16^. These networks coexist in the same cytoplasm and compete for a shared pool of actin monomers^13,17^, yet maintain distinct ABP compositions – fimbrin (Sac6) and capping protein (Cap1/2) localize specifically to branched actin patches, whereas tropomyosin (Tpm1) decorates linear cable arrays^5^. Importantly, while actin disassembly factors are highly conserved between yeast and mammals^12^, the yeast genome encodes only a single gene for each of these factors, enabling precise dissection of their individual contributions without the confounding effects of genetic redundancy.

Here, we use inducible CRISPR interference (CRISPRi) and three-color live-cell imaging to explore the role of actin disassembly factors in promoting proper ABP sorting. We find that Cof1-mediated disassembly is required for maintaining the proper size, function, and ABP composition of cellular actin networks. Specifically, we show that loss of Cof1 activity promotes the assembly of dense, overgrown branched actin networks that are aberrantly decorated by Tpm1. In contrast with prevailing competition models, we show that both branched and linear networks can be co-decorated by Tpm1 and Sac6, and that the assembly of networks with mixed identities does not require formin activity. This work supports a model wherein failure to maintain the proper architecture of actin networks drives mis-localization of network-specific ABPs and severely compromises the functions of these networks.

## RESULTS

### Inducible CRISPRi knockdown of actin disassembly factors

We were interested in exploring the role of five disassembly factors (Srv2, Cof1, Crn1, Twf1, and Aip1) in sorting actin binding proteins (ABPs) to the correct types of actin networks in budding yeast. Null mutants for *SRV2*, *CRN1, TWF1,* and *AIP1* are viable; however, *COF1* is essential for viability^18^. Therefore, we decided to use inducible CRISPR interference (CRISPRi) to knock down the expression of each gene. This approach allowed us to interrogate the function of each gene in a similar manner and within only a few generations (i.e., fewer than 10 divisions) following the induction of transcriptional repression. For each disassembly factor, we designed two guide RNAs (gRNAs; Supplemental Table 1) that were predicted to have the greatest level of transcriptional repression and then cloned each gRNA into a plasmid where their expression was controlled by a tetracycline repressor. This plasmid also contained a yeast centromeric origin of replication and a catalytically inactive Cas9 fused to the MxiI transcriptional repressor (dCas9)^19^.

To test the ability of this system to decrease the expression of our target genes, we individually introduced these plasmids into strains where the target gene was labelled with a C-terminal mNeonGreen (mNG) tag. The only exception was Cof1, where integrating fluorescent tags into the native genomic loci is not possible. Therefore, we integrated an additional copy of Cof1 that contained an internal GFP label^20^. For each strain, we induced knockdown using 250 ng/mL of anhydrotetracycline (ATC) for 16-18 hours and then performed live-cell imaging to quantify the fluorescence intensity of the target gene. We found that at least one gRNA significantly reduced the fluorescence intensity of all five disassembly factors. The strongest reduction in fluorescence intensity for Aip1, Cof1, Crn1, Twf1, and Srv2 was 4.4±0.7, 2.7±0.2, 2.7±0.3, 1.4±0.2, and 3.2±1.0-fold (all reported values represent mean ± 95% CI, unless otherwise indicated; see Supplemental Table 2 for all results from statistical analyses), respectively (Supplemental Figure 1A-B).

### Cofilin is required for proper tropomyosin localization

Next, we asked whether decreasing the expression of actin disassembly factors caused defects in ABP sorting. To test this, we fluorescently labeled the major tropomyosin isoform (mNG-Tpm1) in a strain where actin patches were also labeled (Arc15-mScarlet (mSc)). Tpm1 specifically localizes to linear actin networks (i.e., cables and cytokinetic rings) and does not localize to branched patch networks^21,22^. Therefore, after inducing knockdown of each disassembly factor (Aip1, Cof1, Crn1, Twf1, and Srv2), we used live-cell imaging to quantify how much mNG-Tpm1 was localized to patches. To do this, we used a segmentation strategy to compute the proportion of Tpm1 in patches by dividing the mNG-Tpm1 fluorescence intensity associated with patches by the whole cell fluorescence intensity (Supplemental Figure 2A). In wildtype and uninduced CRISPRi cells this proportion was ∼0.1 (Supplemental Figure 2C, Supplemental Figure 5E), indicating that the majority of mNG-Tpm1 was associated with cables or localized to the cytoplasm. We found that there was no significant change in the proportion of mNG-Tpm1 associated with patches during knockdown of Aip1, Crn1, Twf1, or Srv2 (1.1±0.2, 1.1±0.1, 1.1±0.2, and 1.1±0.1-fold, respectively; Supplemental Figure 2C). However, there was a significant increase in the proportion of mNG-Tpm1 localized to patches during knockdown of Cof1 (3.5±0.3-fold; Figure 1A-B). In addition to recruiting mNG-Tpm1 to their patches, Cof1 mutants were also larger, had a greater number of actin patches, and appeared to have severe defects in cable assembly (e.g., defects in cable length, polarity, and organization). We also noticed that patch networks in Cof1 mutants appear larger than normal and that their shape was highly irregular compared to their usual circular shape.

**Figure 1:**
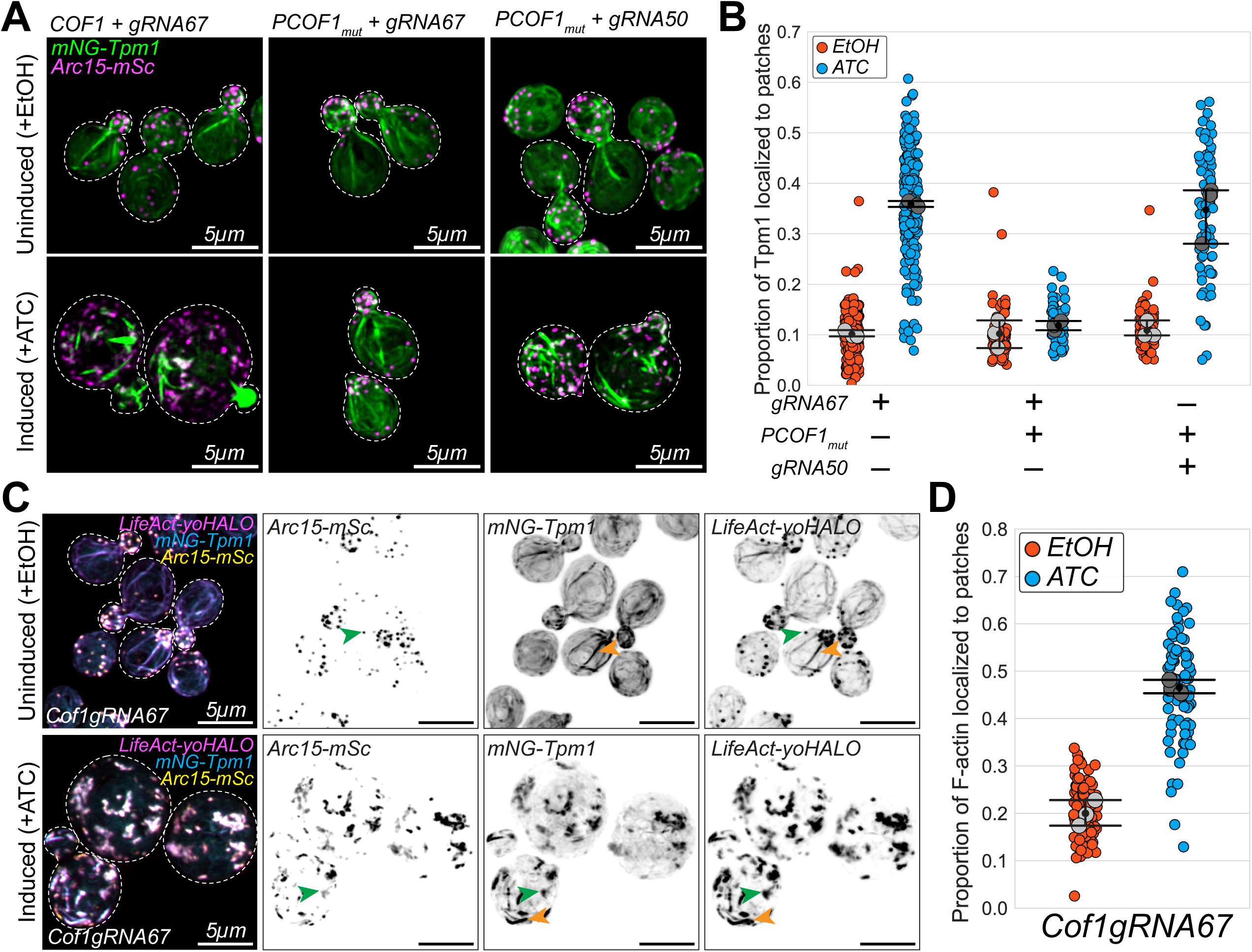
Inducible CRISPRi-mediated knockdown of Cof1 disrupts Tpm1 localization. **(A)** Representative maximum intensity projection images of uninduced (‘EtOH’, top panels) and induced (‘ATC’, bottom panels) Cof1 CRISPRi knockdown mutants that express either Cof1gRNA67 (left and middle panels) or Cof1gRNA50 (right panel) in addition to mNG-Tpm1 and Arc15-mSc. Dashed white line indicates outline of cells traced from brightfield images. All cells express Cof1 from the wildtype promoter. Cells in the middle and right panels express an additional CRISPRi-resistant Cof1 (PCOF1_mut_). Scale bar, 5µm. **(B)** The proportion of mNG-Tpm1 localized to patches in the Cof1 CRISPRi mutants in A. **(C)** Representative maximum intensity projection images of uninduced (‘EtOH’, top panels) and induced (‘ATC’, bottom panels) Cof1gRNA67 CRISPRi knockdown mutants that express LifeAct-yoHALO (magenta), mNG-Tpm1 (cyan), and Arc15-mSc (yellow). Orange arrowheads indicate cable networks. Green arrowheads indicate patch networks. Scale bar, 5µm. **(D)** The proportion of F-actin localized to patches in the Cof1gRNA67 CRISPRi knockdown mutants. In all plots, each colored data point is from a single cell, and larger grey symbols represent the mean from each experiment. Error bar, 95% confidence intervals. Statistical significance determined by students t-test.

To validate these findings, we asked if expression of a CRISPRi-resistant *COF1* could rescue the mNG-Tpm1 mis-localization phenotype. Specifically, we introduced an additional copy of *COF1* whose expression was controlled by a promoter that contained a mutation in the proto-spacer adjacent motif (PAM) site for the gRNA (Cof1gRNA67) used to knock down *COF1* (i.e., the PAM sequence was changed from NGG to NCC). We found that the proportion of mNG-Tpm1 associated with patches in cells expressing CRISPRi-resistant *COF1* (*PCOF1_mut_*) was indistinguishable between uninduced and induced cells (1.2±0.3-fold; Figure 1A-B), indicating rescue of the mNG-Tpm1 mis-localization phenotype. To further validate these results, we induced the expression of a gRNA (Cof1gRNA50) that targets a different PAM site in our CRISPRi-resistant *COF1* strain. We found that expression of this gRNA led to a significant increase in Tpm1 associated with patches (3.2±0.7-fold; Figure 1A-B), recapitulating our earlier measurements using Cof1gRNA67. These findings demonstrate that mis-localization of Tpm1 to patches is the consequence of decreased *COF1* expression and is not likely due to off target CRISPRi gene repression.

Next, we wanted to directly visualize the association of mNG-Tpm1 and Arc15-mSc with F-actin networks. To test this, we generated a three-color live-cell imaging strain that expressed a F-actin reporter (i.e., LifeAct^23,24^) labeled with a codon optimized HALO tag^25^ (LifeAct-yoHALO) in addition to mNG-Tpm1 and Arc15-mSc. We then knocked down Cof1 in this strain and performed live-cell imaging to compare the localization of all three tagged proteins. Immediately prior to imaging, we stained uninduced and induced cells with 5 µM of Janelia Fluor 669 (JF669), a bright, far-red cell permeable dye that covalently binds to yoHALO. In uninduced control cells, we found that actin cables are co-labeled by LifeAct-yoHALO and mNG-Tpm1, while patches are co-labeled by LifeAct-yoHALO and Arc15-mSc (Figure 1C). In contrast, Cof1 mutants had large, misshapen patches that were labeled by all three reporters and unpolarized cables that were co-labeled by LifeAct-yoHALO and mNG-Tpm1 (Figure 1C). To quantify defects in actin homeostasis (i.e., the balance between the assembly of branched and linear networks), we knocked down Cof1 in strains expressing F-actin and patch reporters (LifeAct-mNG and Arc15-mSc, respectively) and then computed the proportion of F-actin localized to patches using segmentation (Supplemental Figure 2A). Consistent with our qualitative observations, we found that Cof1 mutants had a significant increase in the amount of F-actin localized to patches (2.3±0.3-fold; Figure 1D, Supplemental Figure 2E). Taken together, these data indicate that Cof1 activity is required for controlling both the homeostatic balance of actin network assembly, as well as sorting Tpm1 to the correct types of actin networks.

### Myosin-based intracellular transport is disrupted during Cof1 knockdown

In addition to stabilizing actin filaments by inhibiting Cof1 binding^15^, tropomyosin promotes the movement of type V myosins along actin cables^26–28^. Consistent with this role, studies in budding and fission yeast have shown that capping protein mutants, which mis-localize tropomyosin to actin patches, have defects in myosin-mediated intracellular transport, where cargo is diverted from its intended destination^8,9^. Therefore, we were interested in determining whether the Tpm1-decorated patches in Cof1 knockdown cells similarly disrupt myosin-based transport.

First, we added a C-terminal yoHALO tag to Myo2, the type V myosin responsible for intracellular transport in budding yeast^29–31^, and then tracked its localization during Cof1 knockdown. We found that while Myo2-yoHALO was polarized to the bud tip in uninduced control cells, its signal appeared diffuse and unpolarized in Cof1 mutants (Figure 2A). To quantify this, we used line scans to track the fluorescence intensity across the polarity axis of the cell (i.e., from the bud tip to the rear of the mother cell). Consistent with our qualitative assessment, we found that the intensity of Myo2-yoHALO is strongly polarized to the bud tip in uninduced cells, but that the signal is uniform in Cof1 mutants (Figure 2B), suggesting that polarized transport was defective in these cells.

**Figure 2:**
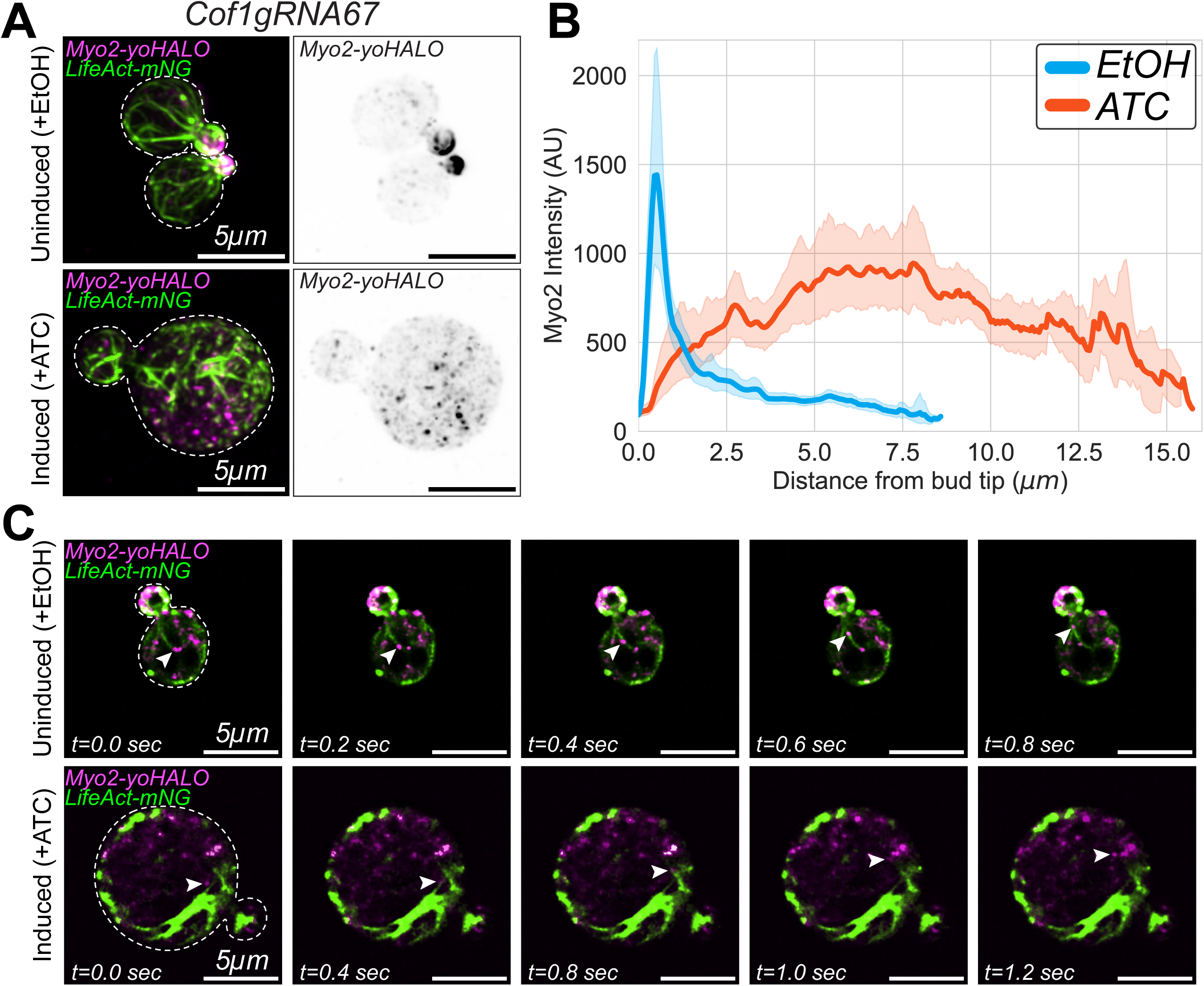
Myosin-based intracellular transport is disrupted in Cof1 mutants. **(A)** Representative maximum intensity projection images of uninduced (‘EtOH’, top panels) and induced (‘ATC’, bottom panels) Cof1gRNA67 CRISPRi knockdown mutants expressing LifeAct-mNG and Myo2-yoHALO. Dashed white line indicates outline of cells traced from brightfield images. Scale bar, 5µm. **(B)** Fluorescent line scan profiles of Myo2-yoHALO along the polarization axis of the cell. **(C)** Representative frames from time series images of Myo2-yoHALO (white arrowheads) moving along actin networks labelled by LifeAct-mNG. Time (seconds) is indicated in the bottom left corner. Dashed white line indicates outline of cells traced from brightfield images. Scale bar, 5µm.

Next, we acquired 2D timelapse images to track the motility of Myo2-yoHALO puncta as they moved along the actin cables labeled by LifeAct-mNG. In control cells, we observed Myo2-yoHALO puncta moving along actin cables and being transported into bud cells (Figure 2C; Supplemental Video 1). In Cof1 knockdown cells, we also observed Myo2-yoHALO puncta moving along cable-like structures; however, due to the severely disorganized actin cytoskeleton in these mutants, they were not transported into bud cells. Instead, the Myo2-yoHALO puncta appear to move along filaments within the disordered F-actin networks until they were deposited at random locations within the mother cell (Figure 2C; Supplemental Video 2). Together, these observations demonstrate that loss of Cof1 activity disrupts polarized intracellular transport, likely because Tpm1 inappropriately decorates branched networks and creates ectopic tracks that misdirect Myo2-dependent cargo.

### Cofilin mediated disassembly is required for proper Tpm1 localization

In addition to its roles in actin filament disassembly (e.g, severing, debranching, and depolymerization), Cof1 can also bind to ADP-actin monomers and inhibit nucleotide exchange^32,33^. In this way, Cof1 can tune the available pool of polymerization competent actin monomers. To determine which of these two functions may contribute to controlling the localization of Tpm1, we tracked the localization of mNG-Tpm1 in strains that express the temperature-sensitive allele *cof1-22*. Prior work has established that *cof1-22* can still bind to G-actin monomers, but has impaired F-actin binding and is defective in filament depolymerization^34,35^. Therefore, we tracked mNG-Tpm1 localization in a strain that expressed the *cof1-22* allele and our patch marker (Arc15-mSc). Compared to cells expressing the *COF1* wildtype allele, we found that *cof1-22* mutants had more mNG-Tpm1 associated with their patches when grown at both the permissive and restrictive temperature (i.e., 2.6±0.6 and 2.0±0.5-fold, respectively; Figure 3A-B). These results agree with our earlier findings using CRISPRi and support a role for Cof1-mediated disassembly in promoting the proper localization of Tpm1.

**Figure 3:**
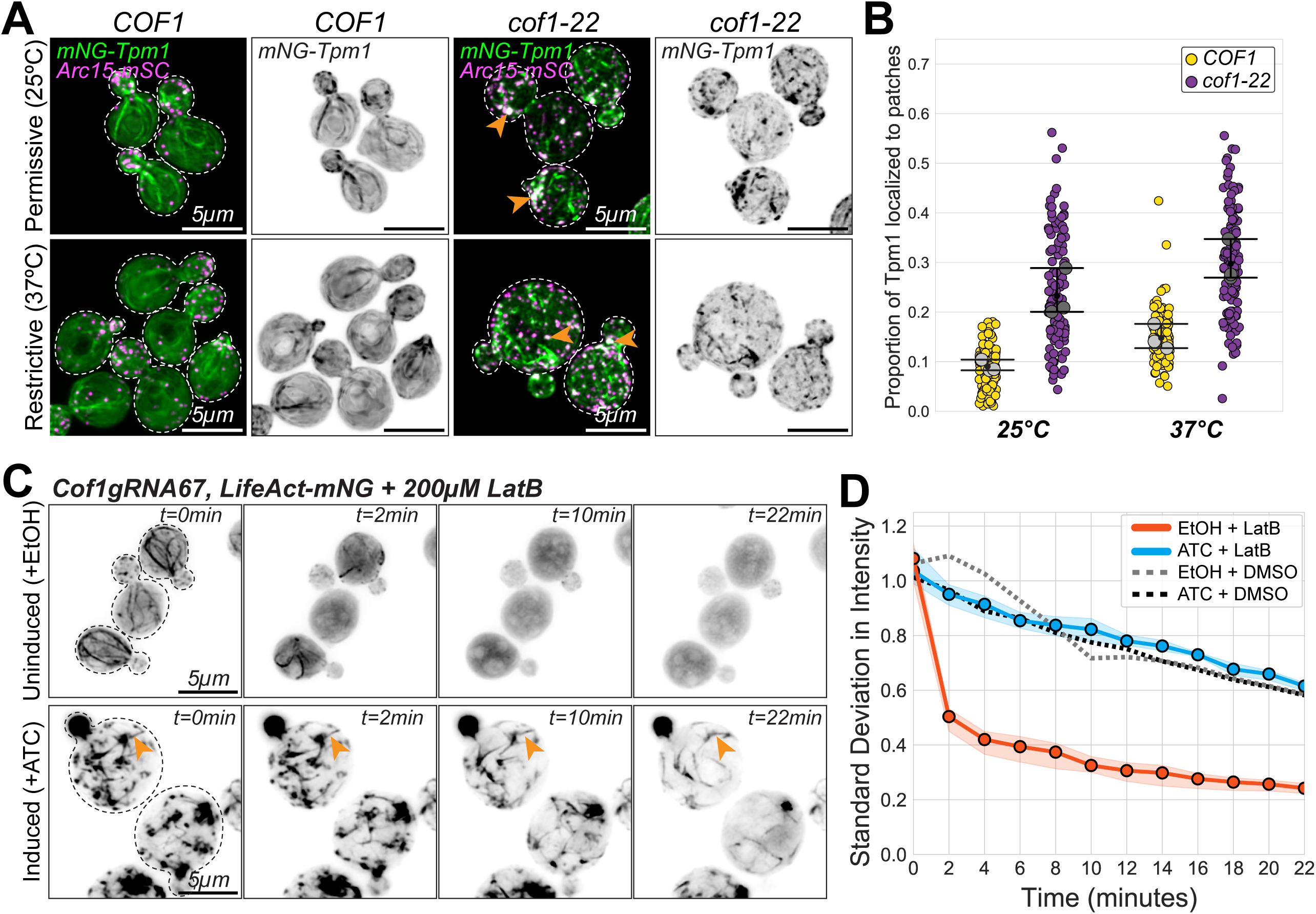
Cof1-mediated disassembly is required for proper Tpm1 localization. **(A)** Representative maximum intensity projection images of cells expressing mNG-Tpm1 and Arc15-mSc, and either COF1 (left panels), or *cof1-22* (right panels) grown at either the permissive (25°C, top panels) or restrictive temperature (37°C, bottom panels). Orange arrowheads indicate patches decorated with mNG-Tpm1. Dashed white line indicates outline of cells traced from brightfield images. Scale bar, 5µm. **(B)** The proportion of mNG-Tpm1 localized to patches in cells from A. Each colored data point is from a single cell, and larger grey symbols represent the mean from each experiment. Error bar, 95% confidence intervals. Statistical significance determined by students t-test. **(C)** Representative maximum intensity projection images of uninduced (‘EtOH’, top panels) and induced (‘ATC’, bottom panels) Cof1gRNA67 CRISPRi knockdown mutants expressing LifeAct-mNG. Cells were treated with 200µM LatB and imaged at the indicated timepoints. Orange arrowheads indicate cable networks that persist after LatB treatment. Scale bar, 5µm. **(D)** The change in LifeAct-mNG fluorescence intensity as a function of time following treatment with 200µM LatB measured from three independent experiments. Solid, colored lines and shading, mean and 95% confidence interval. Dashed lines indicate photobleaching and vehicle controls.

Next, we asked whether network turnover was impaired during Cof1 knockdown. To test this, we treated Cof1 mutants with 200µM Latrunculin B (LatB) to rapidly inhibit actin assembly^36^ and then tracked LifeAct-mNG intensity as actin networks were disassembled. While actin networks in uninduced cells were rapidly disassembled after LatB treatment (i.e., only half of their F-actin structures remained intact after 2 minutes; Figure 3C-D, Supplemental Video 3), networks in Cof1 mutants disassembled very slowly (i.e., after 22 minutes ∼60% of their networks remained intact; Figure 3C-D, Supplemental Video 4). Consistent with prior studies, we noticed that the disassembly rates of different networks were not the same in Cof1 mutants^37^. Specifically, patch networks appeared to disassemble more rapidly than linear networks, and many Cof1 mutants still had intact cable-like structures after 22 minutes of LatB treatment (Figure 3C). These results demonstrate that knockdown of Cof1 reduces the rates of network turnover, likely due to decreased Cof1-mediated disassembly of these networks.

### Actin networks in Cof1 mutants are co-decorated by Tpm1 and Fimbrin (Sac6)

Next, we wanted to determine whether loss of Cof1 activity globally disrupted the localization of network-specific ABPs, or if only Tpm1 localization was disrupted. To test this, we knocked down Cof1 and used three-color live-cell imaging to track the localization of capping protein (Cap2-yoHALO) and fimbrin (Sac6-yoHALO) in strains that also expressed LifeAct-mNG and Arc15-mSc. Normally, neither of these ABPs localize to cables - Cap2 binds the barbed-ends of filaments in patch networks^38^, while Sac6 binds to the sides of branched filaments to bundle them^39,40^. Consistent with this, we found that the localization of Cap2-yoHALO and Sac6-yoHALO in uninduced control cells was restricted to patch networks (Figure 4A-B). After inducing Cof1 knockdown we found that while Cap2-yoHALO localization was still restricted to patches, Sac6-yoHALO appeared to also localize to cable networks (Figure 4A-B). While we did not observe this in all cells or all cables, these data indicate that localization of Sac6 and Tpm1 are similarly disrupted during Cof1 knockdown (Figure 4B-C).

**Figure 4:**
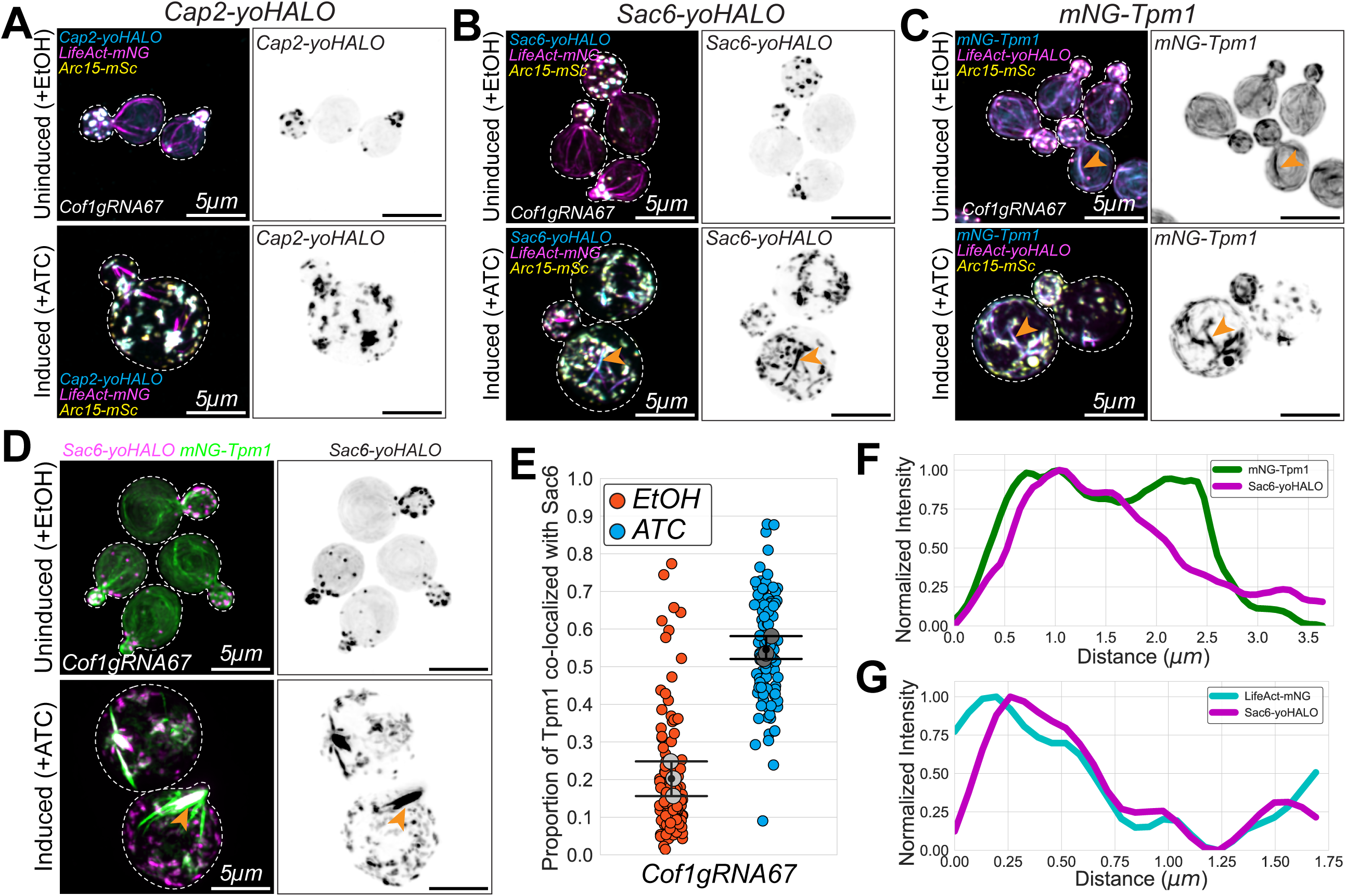
Sac6 and Tpm1 inappropriately colocalize in Cof1 knockdown mutants. **(A-D)** Representative maximum intensity projection images of uninduced (‘EtOH’, top panels) and induced (‘ATC’, bottom panels) Cof1gRNA67 CRISPRi knockdown mutants expressing either: (A) mNGTpm1, LifeAct-mNG, and Arc15-mSc, (B) Cap2-yoHALO, LifeAct-mNG, and Arc15-mSc, (C) Sac6-yoHALO, LifeAct-mNG, and Arc15-mSc, or (D) Sac6-yoHALO and mNG-Tpm1. Orange arrowheads indicate cables co-labelled by LifeAct and Tpm1 (A) or Sac6 (C), or by Sac6 and Tpm1 (D). Dashed white line indicates outline of cells traced from brightfield images. Scale bar, 5µm. **(E)** The proportion of mNG-Tpm1 co-localized with Sac6-yoHALO in cells from D. Each colored data point is from a single cell, and larger grey symbols represent the mean from each experiment. Error bar, 95% confidence intervals. Statistical significance determined by students t-test. **(F-G)** Fluorescent line scan profiles along cables co-labeled by mNG-Tpm1 and Sac6-yoHALO (F) or by LifeAct-mNG and Sac6-yoHALO (G).

These findings were striking, as one of the prevailing models used to explain ABP sorting suggests that tropomyosin and fimbrin directly compete for binding sites on actin filaments, resulting in their exclusive decoration of distinct actin networks^6,7^. Therefore, we were interested in determining whether Sac6 and Tpm1 co-localize during Cof1 knockdown. To test this, we tracked Sac6-yoHALO and mNG-Tpm1 localization during Cof1 knockdown. In uninduced cells, Sac6-yoHALO was localized to patches and not detected on cables, while mNG-Tpm1 was localized to cables and the cytoplasm, as expected (Figure 4D). Upon Cof1 knockdown, however, Sac6-yoHALO and mNG-Tpm1 appeared to co-localize on both patches and cables (Figure 4D). Specifically, there was a 2.7±0.6-fold increase in the amount of mNG-Tpm1 co-localized with Sac6-yoHALO during Cof1 knockdown (Figure 4E). Further, line-scan analysis of individual cables confirmed colocalization Sac6-yoHALO and mNG-Tpm1 on these structures (Figure 4F). Similar results were obtained in strains expressing Sac6-yoHALO and LifeAct-mNG (Figure 4G). Together, these findings indicate that efficient competition between Tpm1 and Sac6 depends on Cof1 activity.

### Tpm1 localization to patches is independent of formin activity

Another model proposes that tropomyosin is preferentially sorted to formin-generated networks due to intrinsic differences in filament architecture that distinguish these filaments from those generated by other nucleators^11,41^. Specifically, experiments in fission yeast demonstrate that tropomyosin isoforms are sorted to distinct networks in a formin-dependent manner, where filaments assembled by specific formins are decorated with specific tropomyosin isoforms^10^. Therefore, we were interested in determining whether Tpm1 mis-localization in Cof1 mutants was due to ectopic formin activity in patches.

First, we tracked formin localization during Cof1 knockdown. To do this, we added C-terminal yoHALO tags to either Bni1 (bud tip localized formin) or Bnr1 (bud neck localized formin) in a strain that expressed reporters of F-actin and patches (LifeAct-mNG and Arc15-mSc, respectively; Figure 5A-B). To compare formin localization during Cof1 knockdown, we used line-scans to generate intensity profiles along the polarization axis of the cell (i.e., from the bud tip to the rear of the mother cell). This analysis showed that both formins had the expected localization in uninduced cells: Bni1-yoHALO signal was focused at the bud tip, and Bnr1-yoHALO was localized to the bud neck (Figure 5C-D)^16,42^. In Cof1 mutants, Bni1 localization was severely disrupted and was no longer polarized to the bud tip (Figure 5B&D). While puncta of Bni1-yoHALO were observed within the mother cell, these did not co-localize with either actin network (Figure 5B). In contrast, Bnr1 continued to localize to the bud neck in many Cof1 mutants (Figure 5A&D). We also observed some mis-localized Bnr1-yoHALO signal in the mother cells of Cof1 mutants, but similar to Bni1, these puncta did not appear to co-localize with either actin network (Figure 5A). Together, these results indicate that formins are not ectopically recruited to patch networks during Cof1 knockdown.

**Figure 5:**
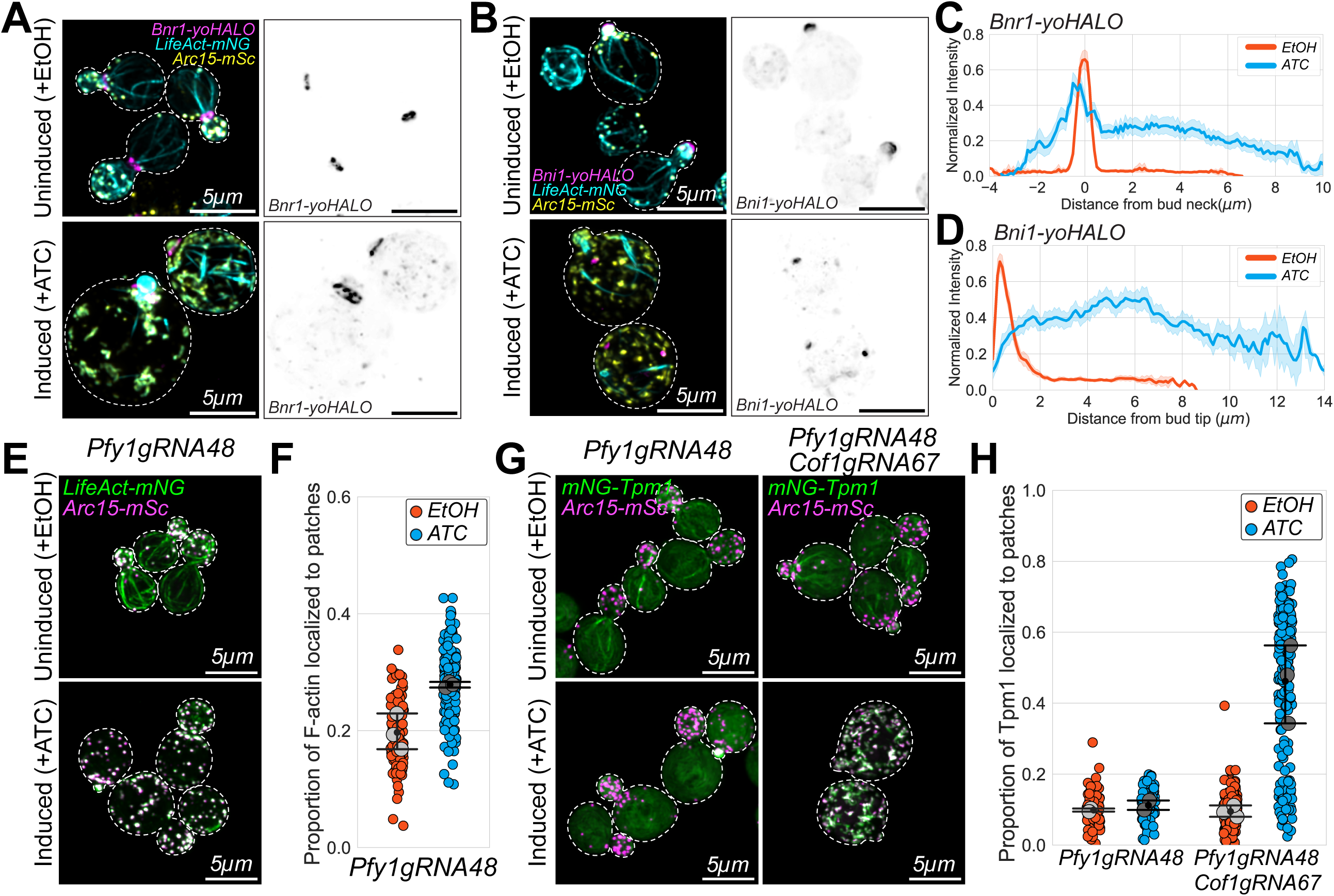
Tpm1 mis-localization to patches is not driven by ectopic formin activity or defects in actin homeostasis. **(A)** Representative maximum intensity projection images of uninduced (‘EtOH’, top panels) and induced (‘ATC’, bottom panels) Cof1gRNA67 CRISPRi knockdown mutants expressing either: (A) Bnr1-yoHALO, LifeAct-mNG, and Arc15-mSc, or (B) Bni1-yoHALO, LifeAct-mNG, and Arc15-mSc. Dashed white line indicates outline of cells traced from brightfield images. Scale bar, 5µm. **(C-D)** Fluorescent line scan profiles of either Bnr1-yoHALO (C) or Bni1-yoHALO (D) along the polarization axis of the cell. **(E)** Representative maximum intensity projection images of uninduced (‘EtOH’, top panels) and induced (‘ATC’, bottom panels) Pfy1gRNA48 CRISPRi knockdown mutants expressing LifeAct-mNG and Arc15-mSc. Scale bar, 5µm. **(F)** The proportion of F-actin localized to patches in the Pfy1gRNA48 CRISPRi knockdown mutants. **(G)** Representative maximum intensity projection images of uninduced (‘EtOH’, top panels) and induced (‘ATC’, bottom panels) Pfy1gRNA48 or Pfy1gRNA48 + Cof1gRNA67 CRISPRi knockdown mutants expressing LifeAct-mNG and Arc15-mSc. Scale bar, 5µm. **(H)** The proportion of mNG-Tpm1 localized to patches in the CRISPRi mutants in G. In all plots, each colored data point is from a single cell, and larger grey symbols represent the mean from each experiment. Error bar, 95% confidence intervals. Statistical significance determined by students t-test.

Although formins were not ectopically recruited to patch networks, it is possible that they contribute to the assembly of Tpm1-decorated patches through transient interactions or by acting during early stages of their assembly. Because deletion of both formins is lethal, we decided to generate a multiplexed CRISPRi knockdown plasmid to simultaneously inhibit expression of Cof1 and profilin (Pfy1), a critical actin regulator that promotes cable assembly by formins^43–45^. Consistent with this role, we found that knockdown of Pfy1 alone globally suppress formin-mediated assembly: there was a significant increase in F-actin localized to patches (1.4±0.2-fold) and many Pfy1 mutants had no cables, although some cells have fewer, shorter cables (Figure 5E-F). Therefore, we next generated a multiplexed CRISPRi plasmid that allowed us to simultaneously knockdown both Cof1 and Pfy1. We found that dual knockdown of Cof1 and Pfy1 did not prevent mis-localization of mNG-Tpm1 to patches; the increase in patch-localized mNG-Tpm1 (4.9 ± 1.4-fold) was comparable to that observed with Cof1 knockdown alone (Figure 5G-H). Moreover, Cof1 knockdown in *bni1Δ* and *bnr1Δ* strains had similar defects in network assembly and organization as cells that expressed both formins (Supplemental Figure 3A-B). Together, these findings indicate that formins do not contribute to the assembly of Tpm1-decorated patches nor do they recruit Tpm1 to patches in Cof1 mutants.

### Tpm1 localization does not depend on maintaining proper actin homeostasis

Next, we asked whether mis-localization of Tpm1 to patches is due to the defects in actin homeostasis. It is possible that Tpm1 localizes to patches in Cof1 mutants simply because these networks are larger than normal and therefore have more potential binding sites. To test this, we induced knockdown of Pfy1 to cause a shift in homeostasis that favored the assembly of patches over cables (Figure 5E-F) and then tracked mNG-Tpm1 localization to patches. We found that there was no difference in the proportion of mNG-Tpm1 associated with patches in Pfy1 mutants (0.11±0.02) when compared to the uninduced control cells (0.10±0.01; Figure 5G-H). Consistent with this, we also found that knockdown of other actin disassembly factors (Aip1, Crn1, and Srv2) caused statistically significant defects in actin network homeostasis that favored patch assembly (e.g., 1.1±0.1, 1.5±0.4, and 1.4±0.1-fold, respectively; Supplemental Figure 2D-E), yet these patches were not decorated by mNG-Tpm1 (Supplemental Figure 2B-C). Overall, these results indicate that mis-localization of Tpm1 to patches is not a consequence of perturbed actin homeostasis.

### Patch number and density increase during Cof1 knockdown

Recent work in budding and fission yeast has shown that capping protein (CP) plays a key role in ABP sorting. In the absence of CP, actin patches become overgrown and are inappropriately co-decorated with tropomyosin, fimbrin, and formins, which disrupts intracellular transport^8,9^. Therefore, we asked whether the ABP-sorting defects observed in Cof1 mutants are similarly driven by patch overgrowth. While our earlier measurements revealed that Cof1 knockdown causes a global shift in the F-actin distribution to favor the assembly of patches over cables (Figure 1D), it is unclear whether this is due to an increase in patch number, size, or density. Therefore, we decided to directly quantify these features of patch architecture in Cof1 mutants and compare these with patches assembled by *cap2Δ* mutants.

First, we deleted *CAP2* from both our F-actin and mNG-Tpm1 reporter strains and found that similar to previous studies there was a significant shift in the F-actin distribution that favored patch assembly (1.9±0.3-fold; Figure 6C, Supplemental Figure 4A) and mNG-Tpm1 was inappropriately recruited to patches (1.8±0.1-fold; Supplemental Figure 4E)^9^. Next, we asked how the number of patches changed in Cof1 and *cap2Δ* mutants. Due to the overlapping signal between neighboring patches, it was difficult to directly count individual patches. Instead, we computed the proportion of cell area that was occupied by patches (i.e., total patch area divided by total cell area) and found that this quantity was significantly increased in both Cof1 and *cap2Δ* mutants. Specifically, there was a 2.7±0.3-fold increase in Cof1 mutants and a 1.4±0.3-fold increase in *cap2Δ* mutants (Supplemental Figure 4B-C).

**Figure 6:**
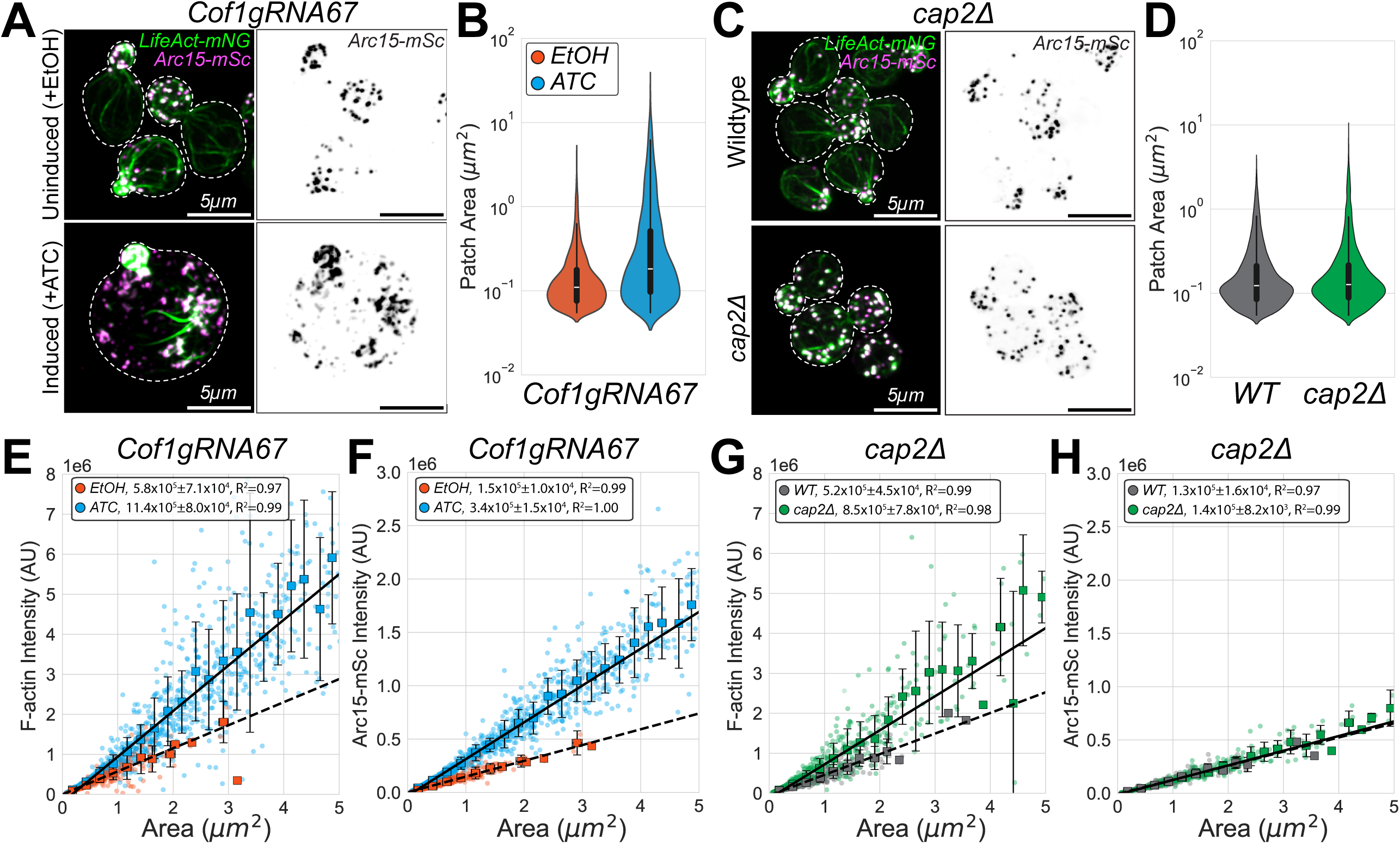
Cof1 mutants assemble overgrown, dense branched actin networks. **(A)** Representative maximum intensity projection images of uninduced (‘EtOH’, top panels) and induced (‘ATC’, bottom panels) Cof1gRNA67 CRISPRi knockdown mutants expressing LifeAct-mNG and Arc15-mSc. Dashed white line indicates outline of cells traced from brightfield images. Scale bar, 5µm. **(B)** Distribution of patch area quantified from uninduced and induced Cof1gRNA67 CRISPRi knockdown mutants in A. **(C)** Representative maximum intensity projection images of wildtype control and *cap2Δ* mutants expressing LifeAct-mNG and Arc15-mSc. Scale bar, 5µm. **(D)** Distribution of patch area quantified from wildtype control and *cap2Δ* mutants in C. **(E-F)** Quantification of patch F-actin density (E) and Arc15 density (F) from Cof1gRNA67 CRISPRi knockdown mutants in A. **(G-H)** Quantification of patch F-actin density (E) and Arc15 density (F) from wildtype control and *cap2Δ* mutants in C. For all plots, each data point represents an individual patch and binned data are represented by large squares. Solid and dashed lines are linear regression fits performed for mutant and control cells, respectively. Error bars, standard deviation.

Next, we compared the sizes of individual patches in these mutants. To do this, we segmented individual patches using our patch reporter (Arc15-mSc) and plotted the distribution of patch area in Cof1 and *cap2Δ* mutants. Using this strategy, we found that while individual patches were 3.9±0.5-fold larger during Cof1 knockdown (Figure 6A-B), patches in *cap2Δ* mutants were the same size as those assembled by wildtype cells (1.2±0.1-fold; Figure 6C-D).

Finally, we compared patch F-actin density in Cof1 and *cap2Δ* mutants. For these analyses, we compared the relationship between LifeAct-mNG fluorescence intensity and patch size. As expected, LifeAct-mNG intensity increased linearly with patch size, indicating that larger patches contain more F-actin. To quantify F-actin density (i.e., LifeAct-mNG intensity per area), we binned these data and measured their slope using linear regression. Through this analysis, we found that F-actin density was 2.0±0.1-fold greater during Cof1 knockdown (Figure 6E) and 1.7±0.1-fold greater in *cap2Δ* mutants (Figure 6G). Taken together, these data indicate that both Cof1 and CP play important roles in maintaining actin homeostasis as well as controlling the density of F-actin in patches, while only Cof1 appears to play a role in controlling the sizes of individual patches.

### The number of filaments in patches increases during Cof1 knockdown

Given that our other homeostasis mutants (i.e., knock down of Pfy1 and other disassembly factors) did not exhibit mis-localization of Tpm1, we decided to focus on understanding how the changes to patch density in Cof1 and *cap2Δ* mutants may promote Tpm1 binding to these networks. Importantly, both Cof1 and CP are thought to control the growth actin filaments in patch networks, albeit via different mechanisms - CP binds to barbed-ends to prevent filament assembly, while Cof1 can disassemble these filaments via severing, depolymerization, or debranching^3,12^. Therefore, we were interested in determining whether Tpm1 mis-localization was driven by defects in the same regulatory process (e.g., controlling filament length) or through distinct processes.

To distinguish between these two possibilities, we decided to quantify the density of Arc15-mSc in patches in Cof1 and *cap2Δ* mutants. Because Arc15 is a subunit of the Arp2/3 complex, it binds to the pointed ends of filaments in patch networks, and its intensity should be proportional to the number of filaments in patches. Therefore, we expected that patches built using longer filaments would have a greater F-actin density, but that their Arc15 density would be unchanged. Alternatively, the density of Arc15 may also increase in denser patches, which would indicate that these patches overgrow due to an increased frequency of branching.

Similar to our quantification of F-actin density, we found that there was a linear relationship between the amount of Arc15-mSc localized to patches and patch size. Using the same linear regression strategy as before, we found that there was 2.3±0.1-fold more Arc15-mSc associated with patches in Cof1 knockdown mutants (Figure 6F), while there was no change (1.0±0.1-fold; Figure 6H) in *cap2Δ* mutants. These data suggest that there are key differences in the patch overgrowth phenotypes exhibited by these mutants: patches in *cap2Δ* mutants are likely composed of overly long filaments, while patches in Cof1 mutants are likely more densely branched. Taken together, these data support a model where controlling the underlying architecture of patch networks plays a critical role in promoting the proper sorting of Tpm1.

## Discussion

Here we used inducible CRISPRi and quantitative live-cell imaging to investigate the role of actin disassembly factors in maintaining proper ABP sorting across distinct actin networks. Our findings reveal that Cof1-mediated disassembly is essential for maintaining the size, function, and ABP composition of actin networks in budding yeast. Specifically, loss of Cof1 activity promotes the assembly of overgrown, disorganized branched actin networks that are inappropriately decorated by Tpm1 and Sac6. A key functional consequence of this mis-sorting is the disruption of myosin-based intracellular transport: Myo2, the type V myosin responsible for polarized cargo delivery, loses its characteristic polarized localization to the bud tip and instead moves along the disordered, Tpm1-decorated patch networks, depositing cargo at ectopic locations within the mother cell. Importantly, these networks with mixed-identities are assembled independently of formin activity and are not simply a consequence of disrupted actin homeostasis. Instead, this work supports a model in which Cof1-mediated control of branched network architecture is a critical determinant of proper ABP sorting.

A central finding of this work is that Cof1 knockdown causes Tpm1 to mis-localize to branched actin patch networks (Figure 1A-B). Among the five disassembly factors we examined, only loss of Cof1 produced this phenotype, even though knockdown of other disassembly factors also caused significant shifts in actin homeostasis (Supplemental Figure 3B-C). This argues against a model in which Tpm1 mis-localizes to patch networks simply because they are larger and therefore have more available binding sites. Consistent with this, knockdown of Pfy1, which also shifts the balance of actin assembly toward patches, did not cause Tpm1 mis-localization (Figure 5E-H). Together, these results indicate that Cof1 plays an important role in maintaining the ABP composition of actin networks that is distinct from its role in regulating actin homeostasis.

Our data further demonstrates that Cof1’s role in ABP sorting is linked to its disassembly activity, rather than its ability to bind G-actin monomers and inhibit nucleotide exchange. The temperature-sensitive *cof1-22* allele, which retains G-actin binding but is impaired in F-actin binding and filament depolymerization^34,35^, phenocopied the Tpm1 mis-localization we observed during CRISPRi knockdown (Figure 3A-B). This supports a model in which Cof1-mediated disassembly (either through severing, depolymerization, or debranching^12^) is required to maintain the proper architecture of branched networks, and that failure to do so results in Tpm1 mis-sorting.

The disruption of Myo2-dependent intracellular transport in Cof1 mutants provides a direct functional readout of ABP sorting failure (Figure 2). In wild-type cells, Tpm1-decorated cables serve as tracks that promote myosin-based transport of growth factors and organelles to the bud^21,29,30^. When Cof1 activity is lost, Tpm1 inappropriately decorates overgrown patch networks, creating ectopic myosin-binding tracks within the mother cell. As a result, Myo2 puncta move along these structures, but fail to reach the bud, yielding a diffuse and unpolarized Myo2 distribution. These observations are consistent with prior work in budding and fission yeast showing that capping protein mutants, which similarly mis-localize tropomyosin to patches, exhibit comparable defects in myosin-mediated cargo transport^8,9^. Together, these findings underscore that the fidelity of ABP sorting is not merely a cytoskeletal organizational feature, but is directly coupled to the execution of essential cellular processes.

A striking finding of this study is that Tpm1 and Sac6 co-decorate actin networks in Cof1 mutants. This directly challenges the prevailing competition model of ABP sorting, which holds that fimbrin and tropomyosin binding to actin filaments is mutually exclusive because they compete for overlapping binding sites^6,7^. This model predicts that the presence of Sac6 on branched patch networks should be sufficient to prevent Tpm1 recruitment. However, our data show that this is not the case in Cof1 mutants, where both proteins simultaneously localize to patches and cables (Figure 4B-G). One possible explanation is that the dense, overgrown patch networks in Cof1 mutants are not fully occupied by Sac6, thereby providing binding sites for Tpm1. Alternatively, studies have demonstrated that cofilin and fimbrin synergize to efficiently displace tropomyosin from actin filaments in vitro^7^. Therefore, loss of Cof1 activity may severely weaken the ability of Sac6 to compete with and displace Tpm1 from filaments. Distinguishing between these possibilities will require future in vitro experiments that directly test how filament architecture and density influence competition between Sac6 and Tpm1.

We also considered whether ectopic formin activity within patch networks could account for Tpm1 mis-localization, given that Tpm1 is normally enriched on formin-generated linear networks^21^. However, several lines of evidence argue against this possibility. Neither Bni1 nor Bnr1 were recruited to patch networks during Cof1 knockdown (Figure 5A-D), and simultaneous knockdown of Cof1 and Pfy1 (which globally suppresses formin-mediated cable assembly) did not prevent Tpm1 mis-localization to patches (Figure 5E-H). Further, Cof1 knockdown in *bni1Δ* and *bnr1Δ* strains produced comparable sorting defects to those observed in cells expressing both formins (Supplemental Figure 3). Collectively, these results demonstrate that Tpm1 recruitment to overgrown patches is likely independent of formin activity and is therefore not driven by an intrinsic sorting preference of Tpm1 for formin-assembled filaments.

Comparisons between Cof1 and CP mutants provides additional mechanistic insight into how branched network architecture controls ABP sorting. Both Cof1 knockdown and *cap2Δ* increased F-actin density within patches and caused Tpm1 mis-localization, yet the underlying architectural changes differed between these mutants (Figure 6, Supplemental Figure 4). In Cof1 mutants, individual patches were significantly larger and contained more Arp2/3 complex, likely indicating increased branching frequency. In contrast, individual patches in *cap2Δ* mutants were not significantly larger, and contained similar levels of Arp2/3 complex, suggesting that longer filaments, rather than more frequent branching, account for the increased density in these cells. This indicates that while distinct changes in patch architecture can result in Tpm1 mis-localization, elevated F-actin density within patches may be a shared feature that underlies sorting defects, regardless of the specific mechanism by which it arises (i.e., changes in filament length or branching frequency). The specific increase in Arp2/3 density observed in Cof1 mutants further suggests that elevated branching may be particularly disruptive to normal ABP sorting, perhaps by generating a more complex architecture that cannot be efficiently saturated by Sac6. Importantly, the functional outcomes in both mutants are similar (i.e., Tpm1 mis-sorting and disrupted myosin-based transport) highlighting that the architectural integrity of branched networks, rather than any single molecular defect, is the key upstream determinant of transport fidelity.

Taken together, our findings support a model for ABP sorting in which Cof1-mediated disassembly maintains the proper architecture of branched actin networks, and that this is required to maintain the competitive interactions that normally partition network-specific ABPs to distinct structures. In this model, when disassembly fails, networks overgrow beyond an architectural regime in which competitive sorting between Sac6 and Tpm1 can effectively operate, enabling the promiscuous association of normally network-specific ABPs. The resulting networks with mixed identities compromise downstream cellular functions: cargo undergoing intracellular transport is diverted from its intended destination because Tpm1-decorated patch networks create aberrant, non-polarized myosin tracks.

This view of ABP sorting complements and extends current competition models by identifying network architecture as a key upstream determinant of sorting fidelity, and highlights actin disassembly as an active regulator of cytoskeletal identity. Future work will be needed to determine how broadly this principle applies, particularly in cell types that express a more complex array of disassembly factors that may provide additional layers of regulation, but where the core relationship between network architecture and ABP sorting is likely to be conserved.

## Methods and Materials

### Plasmids and yeast strains

All yeast strains used in this study are in the W303 background. A complete list of strains used is available in Supplemental Table 3 and a complete list of plasmids used is available in Supplemental Table 4. Strains expressing C-terminal tags at endogenous loci were generated by PCR-based homology repair^46^ by lithium acetate/polyethylene glycol transformation with PCR products amplified from pFA6A vectors using primers with 40-bp homology arms as described. Transformants that grew on selective medium were screened for expression of the tagged protein, and then correct integration into the genome was confirmed by PCR analysis.

Guide RNA sequences were designed using the “Yeast CRISPRi guide selection and evaluation” tool (https://lp2.github.io/yeast-crispri/)^19^. For each gene, we selected the top two gRNA sequences that had Quality Score of 2 and binding sites within 150 bp upstream of the transcription start site. For each gRNA, we ordered a synthesized double stranded DNA oligo (IDT gBlocks) that contained the gRNA sequence flanked by 60 bp regions that were homologous to the insertion site in the plasmid. To clone gRNA containing gBlocks into the inducible CRISPRi plasmid (*pRS416-dCas9-Mxi1-URA + TetR + pRPR1(TetO)-NotI-gRNA*; Addgene plasmid # 73796), we digested the vector with NotI-HF (New England Biolabs) overnight and use homology-based cloning (GeneArt Gibson Assembly HiFi Master Mix, ThermoFisher Scientific) to assemble the final plasmid. Insertion of the gRNA was confirmed via Sanger sequencing (GENEWIZ) and validated plasmids were then introduced into yeast strains via lithium acetate/polyethylene glycol transformation. Transformants were selected using selective media. To express two gRNAs, we first individually cloned each gRNA as described above. Then, we used a universal primer set was used to amplify the entire inducible gRNA cassette from one plasmid and insert it using homology-based cloning (GeneArt Gibson Assembly HiFi Master Mix, ThermoFisher Scientific) into a *SacII* site in a plasmid containing the second gRNA. The presence of both gRNAs was validated using Sanger sequencing (GENEWIZ) and whole plasmid sequencing (Plasmidsaurus) before dual CRISPRi mutants were generated as described above.

To generate a CRISPRi-resistant *COF1,* we PCR amplified *COF1* from genomic DNA and inserted this using homology-based cloning into an integrating plasmid containing the *COF1* promoter sequence. We then ordered a double stranded DNA oligo (IDT gBlocks) were we had changed the PAM sequence for Cof1gRNA67 from NGG to NCC. We then used homology-based cloning (GeneArt Gibson Assembly HiFi Master Mix, ThermoFisher Scientific) to integrate this mutated promoter region into the integrating plasmid containing *COF1* and confirmed proper insertion and presence of the mutated PAM site via Sanger sequencing (GENEWIZ). We then linearized this plasmid via PCR and integrated it into the *TRP* locus as an additional copy in the appropriate yeast strains. Following confirmation using PCR, we then introduced the Cof1gRNA67 and Cof1gRNA50 CRISPRi plasmids as described above.

Generation of the yoHALO pFA6a plasmid was achieved by PCR amplification of yoHALO from *YIplac211-Sec31-yoHalo* (Addgene plasmid # 115423) and homology-based cloning (GeneArt Gibson Assembly HiFi Master Mix, ThermoFisher Scientific) to replace mNeonGreen from our *pFA6a-mNeonGreen::URA3* and *pFA6a-mNeonGreen::HIS3* plasmids. Transformants were screened for growth on selective media and confirmed via Sanger sequencing (GENEWIZ) and whole plasmid sequencing (Plasmidsaurus).

### Induction of gRNA expression

To induce expression of the gRNA for each CRISPRi knockdown strain, single colonies were grown overnight in synthetic media lacking either uracil or tryptophan at 25°C. After overnight growth, ∼1-2µL of overnight culture was diluted into 5mL of fresh media supplemented with 6.25µL of either 100% ethanol (EtOH) or 0.2mg/mL anhydrotetracycline (ATC; 250ng/mL final concentartion). Cultures were then grown for 16-18 hours at 25°C before preparation for live-cell imaging.

### Live cell imaging

Cells were grown at 25°C to mid-log phase (OD600 ∼0.3-0.5) in yeast synthetic complete media (SCM), or as described above for inducing gRNA expression. Cells were concentrated via centrifugation (8000xg, 1min) in low-retention 1.7mL Eppendorf tubes and 8µL of concentrated cell suspension was added to a 1.2% agarose pad made with the appropriate media supplemented with EtOH or ATC. Three-dimensional (3D) stacks were acquired at 0.2 μm intervals on a Nikon Ti2E inverted confocal microscope equipped with a CSU-W1 spinning-disk head (Yokogawa, Tokyo, Japan) and a Prime BSI sCMOS camera (Teledyne Photometrics) controlled by Nikon Imaging Software (NIS) - Elements Advanced Research software using a 100X, 1.45 NA objective. 3D stacks were acquired for the entire height of the cell. Images were processed using denoising and deconvolution algorithms using NIS-Elements Advanced Research software (Nikon). Following processing, images were subjected to a customized image processing pipeline using Python scripts. Briefly, this pipeline generated sum and maximum intensity projections of each channel before subtracting background fluorescence intensity and generating segmentation masks for actin patches (using Arc15-mSc signal) or the entire cell (using brightfield images). Segmentation of Arc15-mSc images was performed using a manually determined threshold value, while brightfield images were segmented using YeastSpotter^47^. Following the generation of projection images and segmentation masks, individual cells were selected for analysis using custom ImageJ macros that quantified the fluorescence intensity within the entire cell and within patch regions. Further analyses and plotting of these data were performed using custom Python scripts.

### HALO staining

Cells expressing yoHALO tagged proteins were grown as described above. Cells were concentrated via centrifugation (8000xg, 1min) in low-retention tubes and stained with 5µM of Janelia Fluor 669 (JF669) for 30 minutes at 25 °C while protected from ambient light. Following staining, cells were washed with the appropriate media three times and then mounted for imaging on a 1.2% agarose pad and covered with a cleaned coverslip. Coverslips were submerged in 2% Micro-90 and cleaned using a sonicating water bath for 1 hour, followed by washes in 70% and 100% EtOH for 1 hour each.

### Myo2 time series imaging

Cells were grown, stained, and mounted onto 1.2% agarose pads as described above. Time series images were acquired at 5 frames per second using triggered acquisition to allow for rapid imaging of 488 and 561 channels. Images were processed as described above using NIS-Elements Advanced Research software (Nikon),

### *cof1-22* induction

Cells were grown at 25°C to mid-log phase (OD600 ∼0.3-0.5) in yeast synthetic complete media (SCM). Cultures were then shifted to 37°C for 1 hour before being concentrated into low-retention 1.7mL Eppendorf tubes and mounted onto 1.2% agarose pads as described above. During imaging, the appropriate temperature (either 25°C or 37°C) was maintained using a cage incubator (Okolab).

### Latrunculin B (LatB) treatment

Cells were grown at 25°C to mid-log phase (OD600 ∼0.3-0.5) and concentrated into low-retention 1.7mL Eppendorf tubes as described above for inducing gRNA expression. Cells were then added to individual wells of a glass-bottom 96-well imaging plate coated with 2mg/mL Concanavalin A (ConA). Cells were allowed to adhere to the bottom of the well until the desired density was achieved. Following this, the cell suspension was aspirated and replaced with fresh media to prevent further cell adhesion. Immediately before acquisition, all media was removed from the well and fresh media containing either 200µM LatB or DMSO was added and time series imaging was started. 3D stacks were acquired for the entire height of the cell at 0.2 μm intervals with a 2 minute time delay for 22 minutes total. Images were processed as described above using NIS-Elements Advanced Research software (Nikon), and custom ImageJ macros were used to track how the intensity changed over time. For this analysis, we found that the standard deviation in intensity was more closely correlated with the presence of intact actin networks than mean or total intensity, as there was greater variability in signal intensity (bright structures with dim cytosolic signal) when cables and patches were intact compared to the uniform cytosolic intensity when no actin structures were present.

### Formin localization

Images were acquired and processed as described above. ImageJ was used to perform linescan analyses on these images, where we generated a fluorescence profile along three defined points for each cell: the bud tip, bud neck, and rear of the mother cell. We then used custom Python scripts to align these profiles so that all Bni1-yoHALO profiles originate from the bud tip, while all Bnr1-yoHALO are aligned at the bud neck.

### Patch area and density quantification

Segmentation masks of patches and background subtracted sum intensity projections of live-cell images were generated as described above using custom Python scripts. Custom ImageJ macros were used to quantify patch area and intensity measurements for the appropriate channels (either LifeAct-mNG or Arc15-mSc). These data were then analyzed using custom Python scripts that binned data based on patch area, using 21 evenly spaced bins between the smallest measured patch in control cells and 5µm^2^. For each bin, the mean and 95% confidence intervals were determined and plotted in addition to the raw data. Linear regression was performed using first 9 bins (i.e., all bins smaller than ∼2µm^2^) as bins in control cells rarely had more than one patch larger than 2µm^2^.

### Statistical analysis

All experiments were repeated using three independent biological replicates. Summary statistics (mean, 95% confidence intervals) were determined using Python. Statistical significance was determined by Student’s t test. Differences were considered significant if p≤0.05.

## Data availability

All images and data are archived at Zenodo, and source code is available at GitHub.

## Acknowledgements

We thank the Lavis lab and Open Chemistry team (Janelia) for the gift of JF and JFX dyes. The inducible CRISPRi plasmid was a gift from Ronald Davis (Addgene plasmid # 73796; http://n2t.net/addgene:73796; RRID:Addgene_73796). The yoHALO containing plasmid (YIplac211-Sec31-yoHalo) was a gift from Benjamin Glick (Addgene plasmid # 115423; http://n2t.net/addgene:115423; RRID:Addgene_115423).

---

**Supplemental Figure 1:**
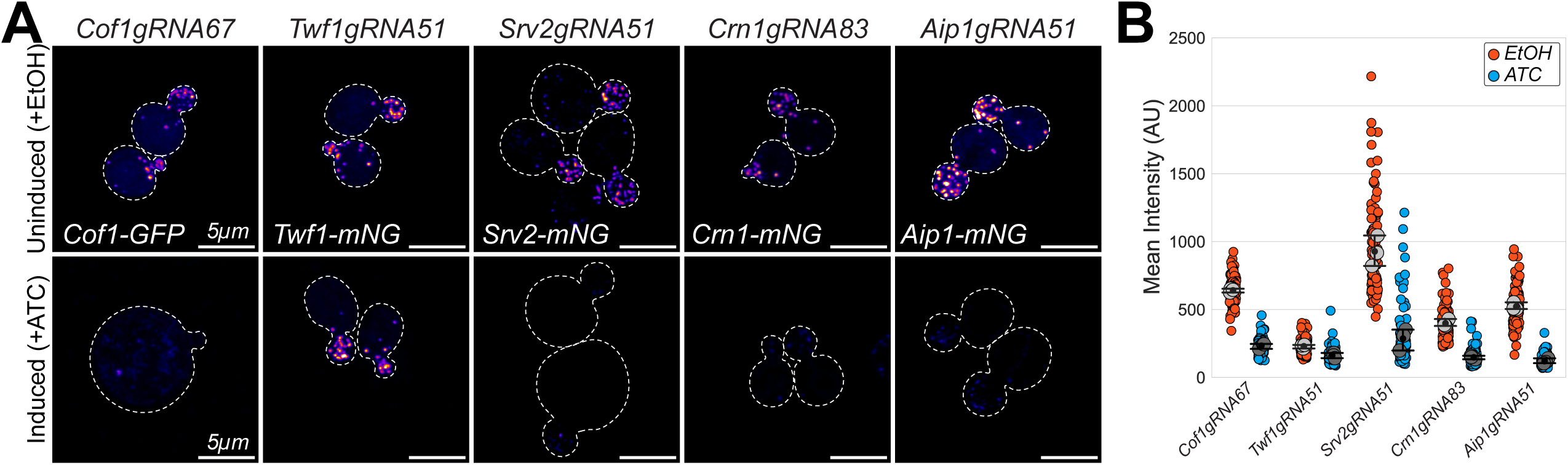
Inducible CRISPRi knockdown of actin disassembly factors. **(A)** Representative maximum intensity projection images of uninduced (‘EtOH’, top panels) and induced (‘ATC’, bottom panels) CRISPRi knockdown mutants expressing the target gene labelled with either mNeonGreen (mNG) or GFP. Dashed white line indicates outline of cells traced from brightfield images. Scale bar, 5µm. **(B)** The mean fluorescence intensity of each target gene in the CRISPRi mutants from A. Each colored data point is from a single cell, and larger grey symbols represent the mean from each experiment. Error bar, 95% confidence intervals. Statistical significance determined by students t-test.

**Supplemental Figure 2:**
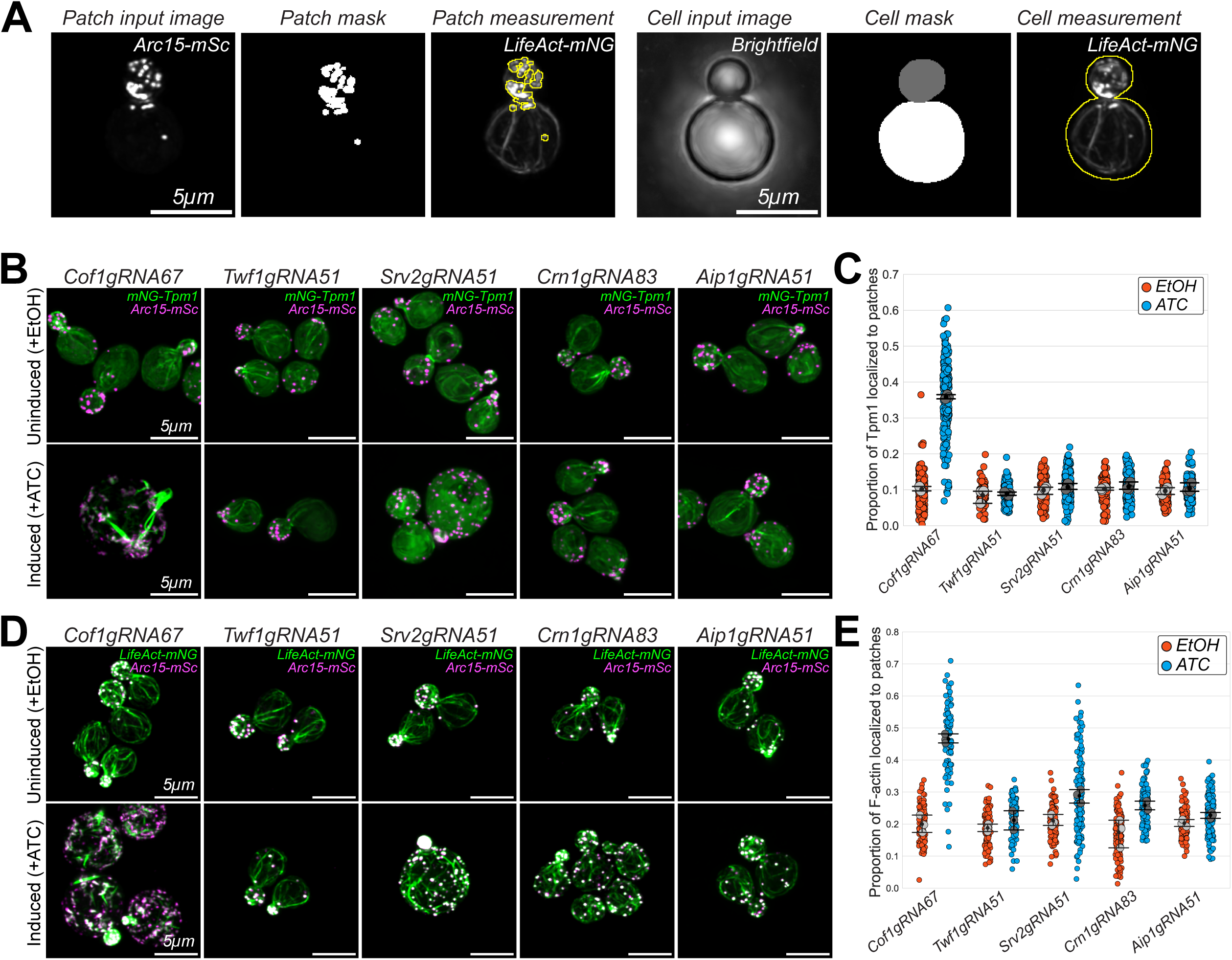
Quantification of changes in mNG-Tpm1 and LifeAct-mNG intensity associated with actin patches in actin disassembly CRISPRi mutants. **(A)** Schematic of the segmentation strategy used to quantify the fluorescence intensity of either mNG-Tpm1 or LifeAct-mNG associated with patches in CRISPRi mutants. Maximum intensity projection images of Arc15-mSc (‘Patch input image’) were used to generate segmentation masks (‘Patch Mask’). The mask was then applied to the corresponding LifeAct-mNG (‘Patch measurement’) or mNG-Tpm1 channel from the same cell to quantify the total fluorescence intensity of either reporter associated with patches. From the same cell, we used a brightfield image (‘Cell input image’) to segment the entire cell (‘Cell mask’) using YeastSpotter. This mask was then applied to the corresponding LifeAct-mNG (‘Cell measurement’) or mNG-Tpm1 channel from the same cell to quantify the total fluorescence intensity in the cell. Scale bar, 5µm. **(B)** Representative maximum intensity projection images of uninduced (‘EtOH’, top panels) and induced (‘ATC’, bottom panels) actin disassembly CRISPRi knockdown mutants that express mNG-Tpm1 and Arc15-mSc. The target gene and PAM site for each gRNA is indicated above each image. Dashed white line indicates outline of cells traced from brightfield images. Scale bar, 5µm. **(C)** The proportion of mNG-Tpm1 localized to patches in the CRISPRi mutants in B. **(D)** Representative maximum intensity projection images of uninduced (‘EtOH’, top panels) and induced (‘ATC’, bottom panels) actin disassembly CRISPRi knockdown mutants that express LifeAct-mNG and Arc15-mSc. The target gene and PAM site for each gRNA is indicated above each image. Dashed white line indicates outline of cells traced from brightfield images. Scale bar, 5µm. **(C)** The proportion of F-actin localized to patches in the CRISPRi mutants in D. For all plots, each colored data point is from a single cell, and larger grey symbols represent the mean from each experiment. Error bar, 95% confidence intervals. Statistical significance determined by students t-test.

**Supplemental Figure 3:**
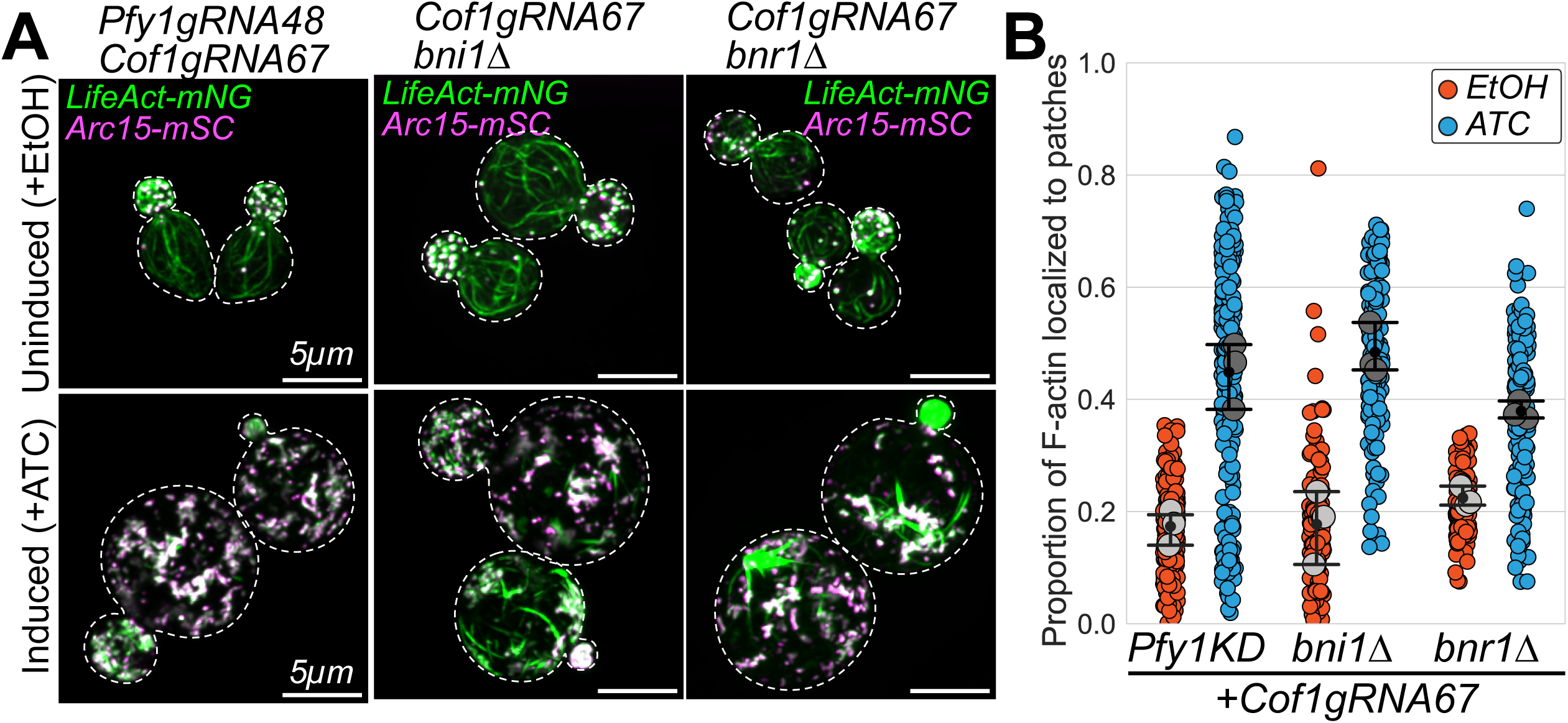
Formin mutants have similar actin homeostasis defects as cells that express both formins. **(A)** Representative maximum intensity projection images of uninduced (‘EtOH’, top panels) and induced (‘ATC’, bottom panels) CRISPRi knockdown mutants that express Pfy1gRNA48 + Cof1gRNA67 (left), or Cof1gRNA67 in *bni1Δ* (center) or *bnr1Δ* (right). All cells also express LifeActNG and Arc15-mSc. Dashed white line indicates outline of cells traced from brightfield images. Scale bar, 5µm. **(B)** The proportion of F-actin localized to patches in the mutants in A.

**Supplemental Figure 4:**
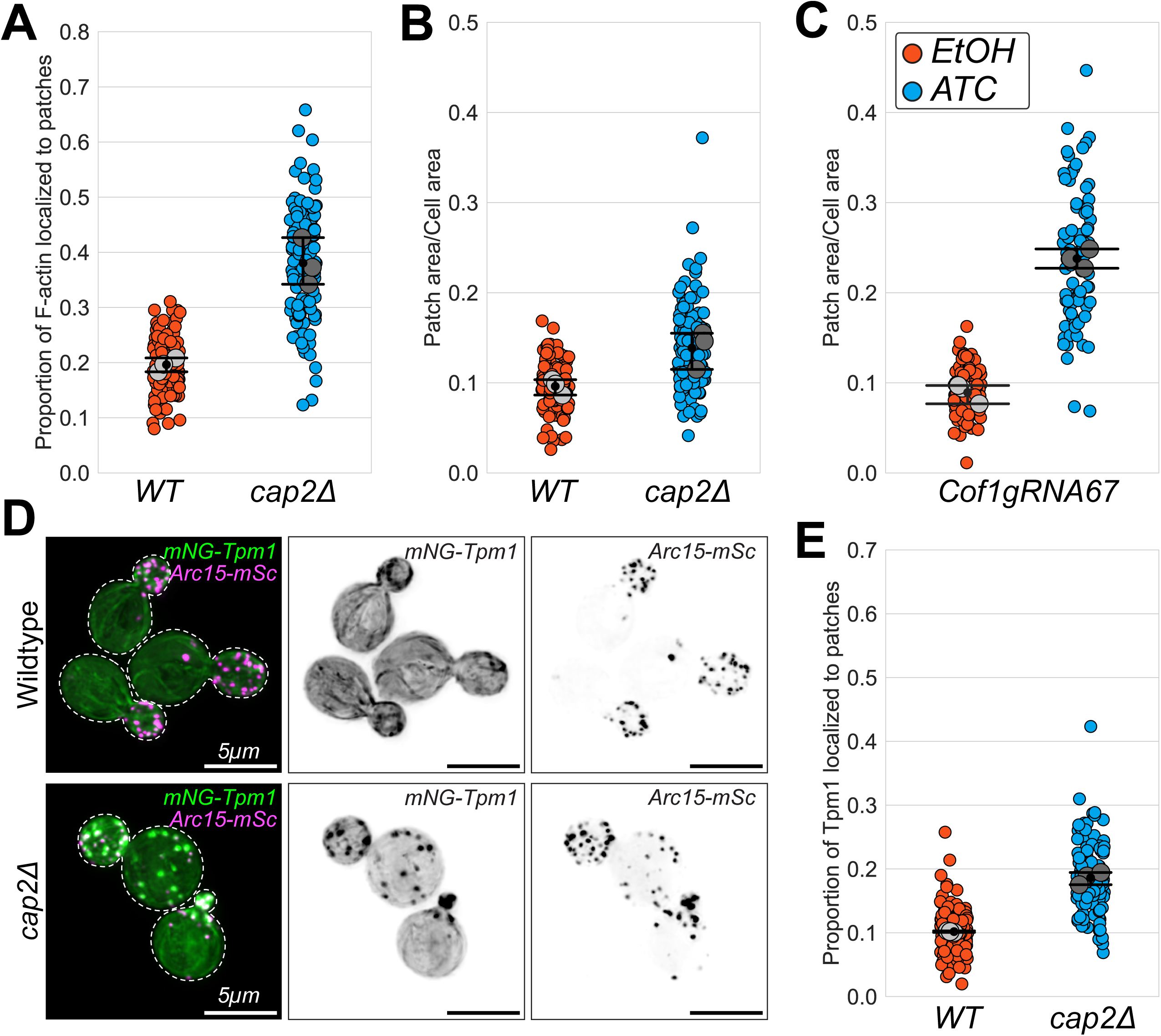
Quantification of actin patch defects in Cof1 CRISPRi and *cap2Δ* mutants. **(A)** The proportion of F-actin localized to patches in *cap2Δ* mutants in Figure 5C. **(B-C)** The proportion of cell area occupied by patches in *cap2Δ* mutants (B) and Cof1gRNA67 mutants (C) from Figure 5A&C. **(D)** Representative maximum intensity projection images of wildtype control and *cap2Δ* mutants expressing mNG-Tpm1 and Arc15-mSc. Scale bar, 5µm. **(E)** The proportion of mNG-Tpm1 localized to patches in *cap2Δ* mutants in D. For all plots, each colored data point is from a single cell, and larger grey symbols represent the mean from each experiment. Error bar, 95% confidence intervals. Statistical significance determined by students t-test.

**Supplemental Movie 1:** Representative 2D time series movie of Myo2-yoHALO (white arrowheads) moving along actin networks labelled by LifeAct-mNG in uninduced Cof1gRNA67 CRISPRi mutants. Video is played at 5 frame per second. Time (seconds) is indicated in the top right corner. Dashed white line indicates outline of cells traced from brightfield images. Scale bar, 5µm.

**Supplemental Movie 2:** Representative 2D time series movie of Myo2-yoHALO (white arrowheads) moving along actin networks labelled by LifeAct-mNG in induced Cof1gRNA67 CRISPRi mutants. Video is played at 5 frame per second. Time (seconds) is indicated in the top right corner. Dashed white line indicates outline of cells traced from brightfield images. Scale bar, 5µm.

**Supplemental Movie 3:** Representative maximum intensity projections from 3D time series images of uninduced Cof1gRNA67 CRISPRi mutants expressing LifeAct-mNG treated with 200µM LatB. Video starts immediately after LatB treatment begins and is played at 1 frame per second. Time (minutes) is indicated in the top right corner. Scale bar, 5µm.

**Supplemental Movie 4:** Representative maximum intensity projections from 3D time series images of induced Cof1gRNA67 CRISPRi mutants expressing LifeAct-mNG treated with 200µM LatB. Video starts immediately after LatB treatment begins and is played at 1 frame per second. Time (minutes) is indicated in the top right corner. Scale bar, 5µm.

**Supplemental Table 1:**
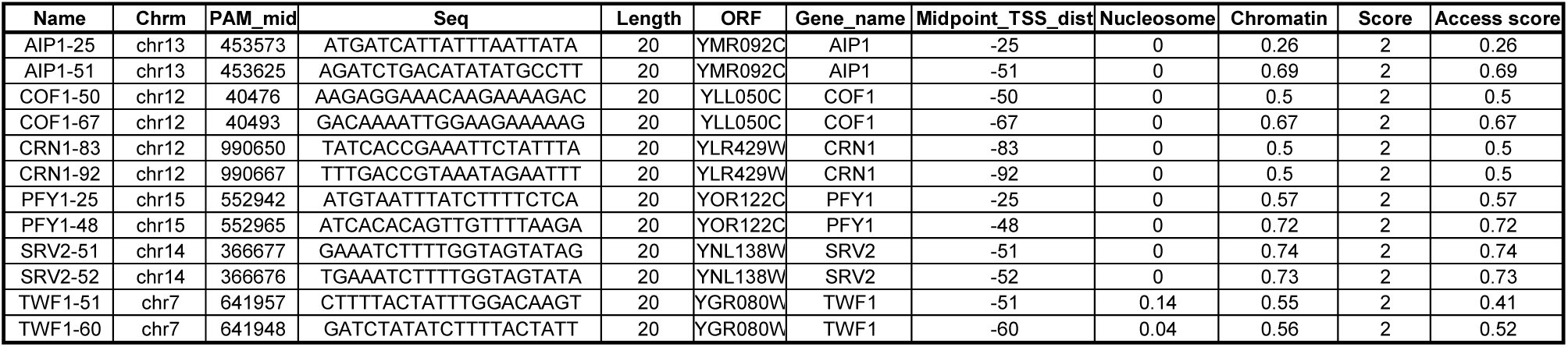
List of guide RNAs (gRNAs) used in this study.

**Supplemental Table 2:**
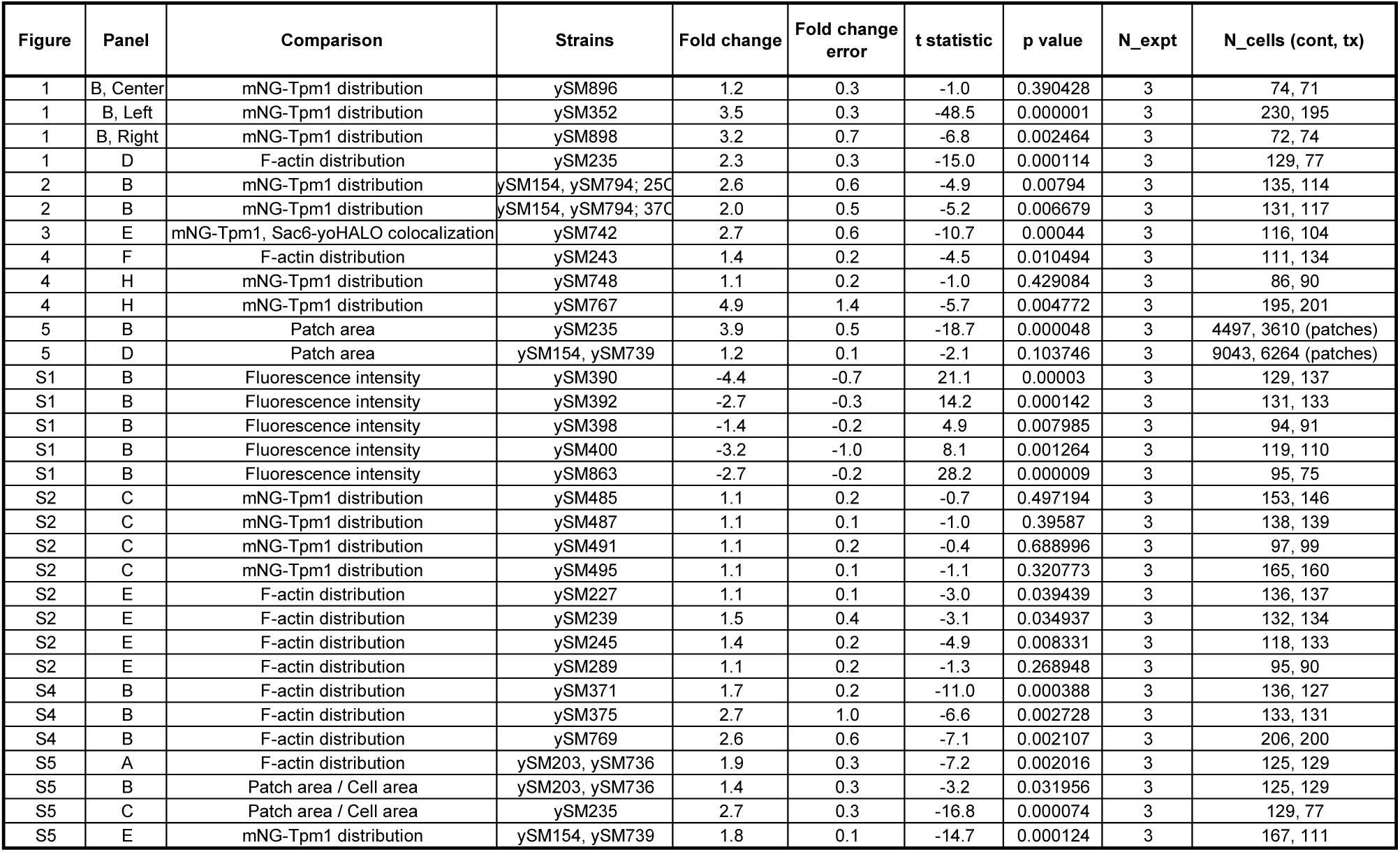
Summary of statistical comparisons made in this study.

**Supplemental Table 3:**
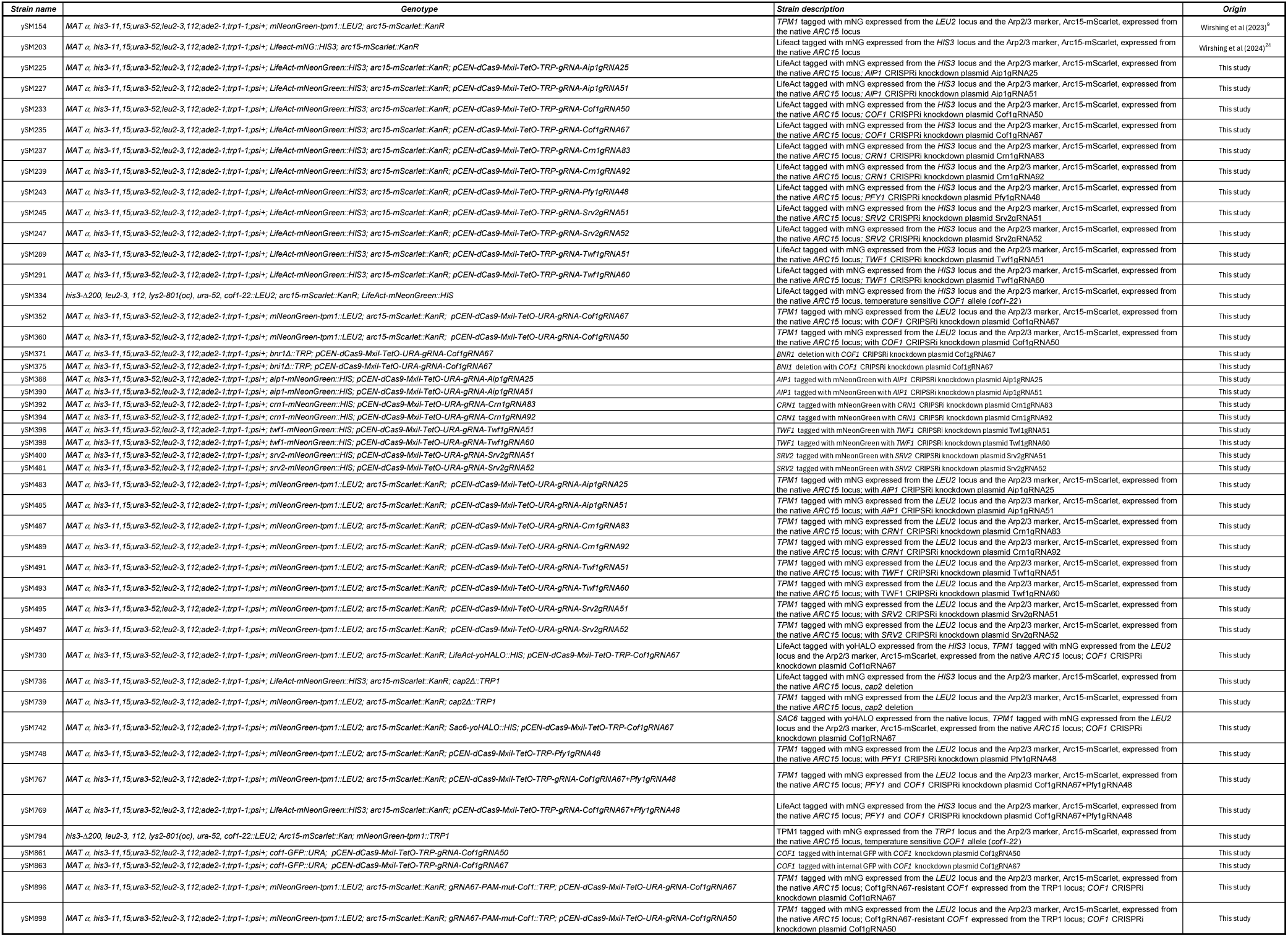
List and description of yeast strains used in this study.

**Supplemental Table 4:**
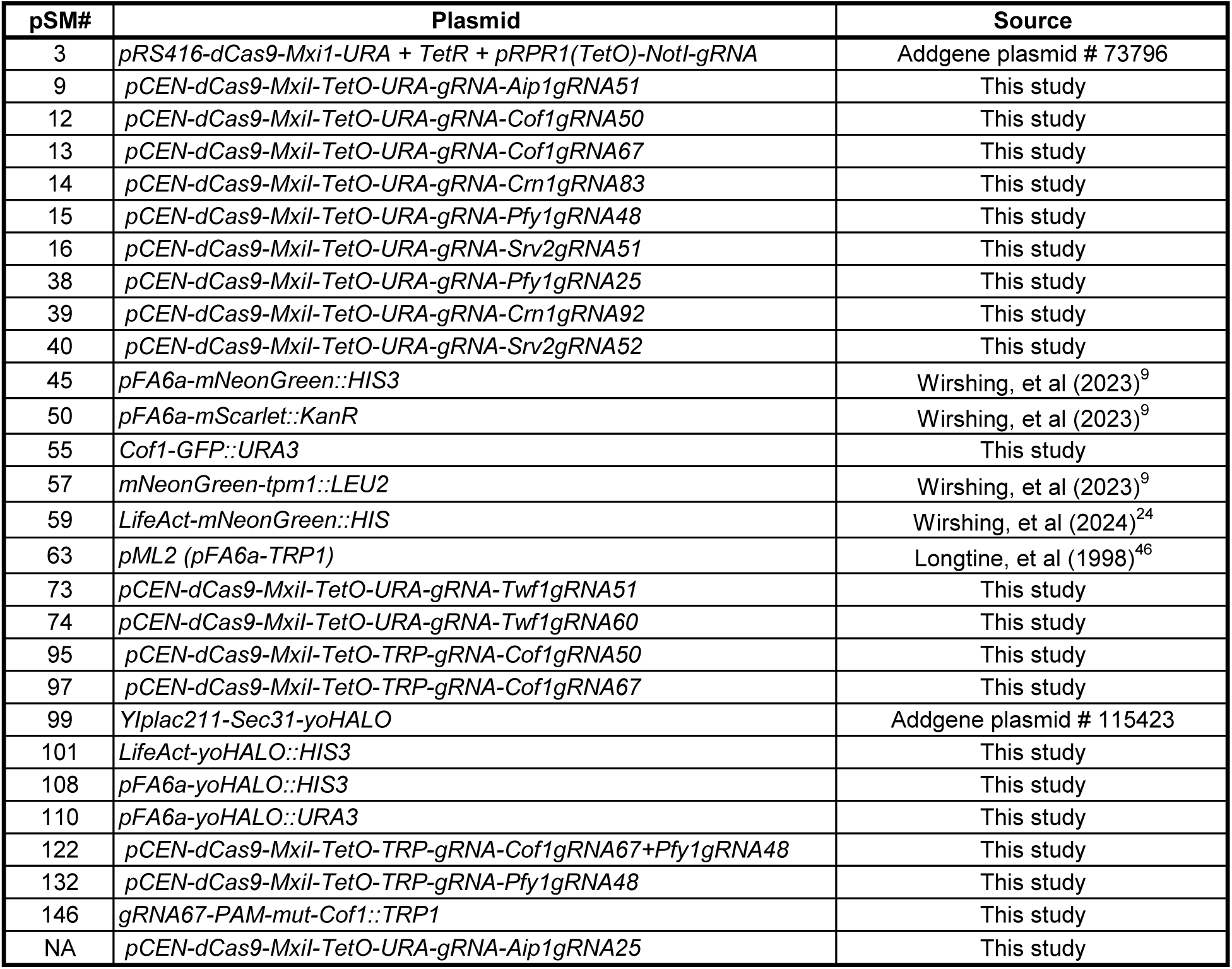
List of plasmids used in this study.

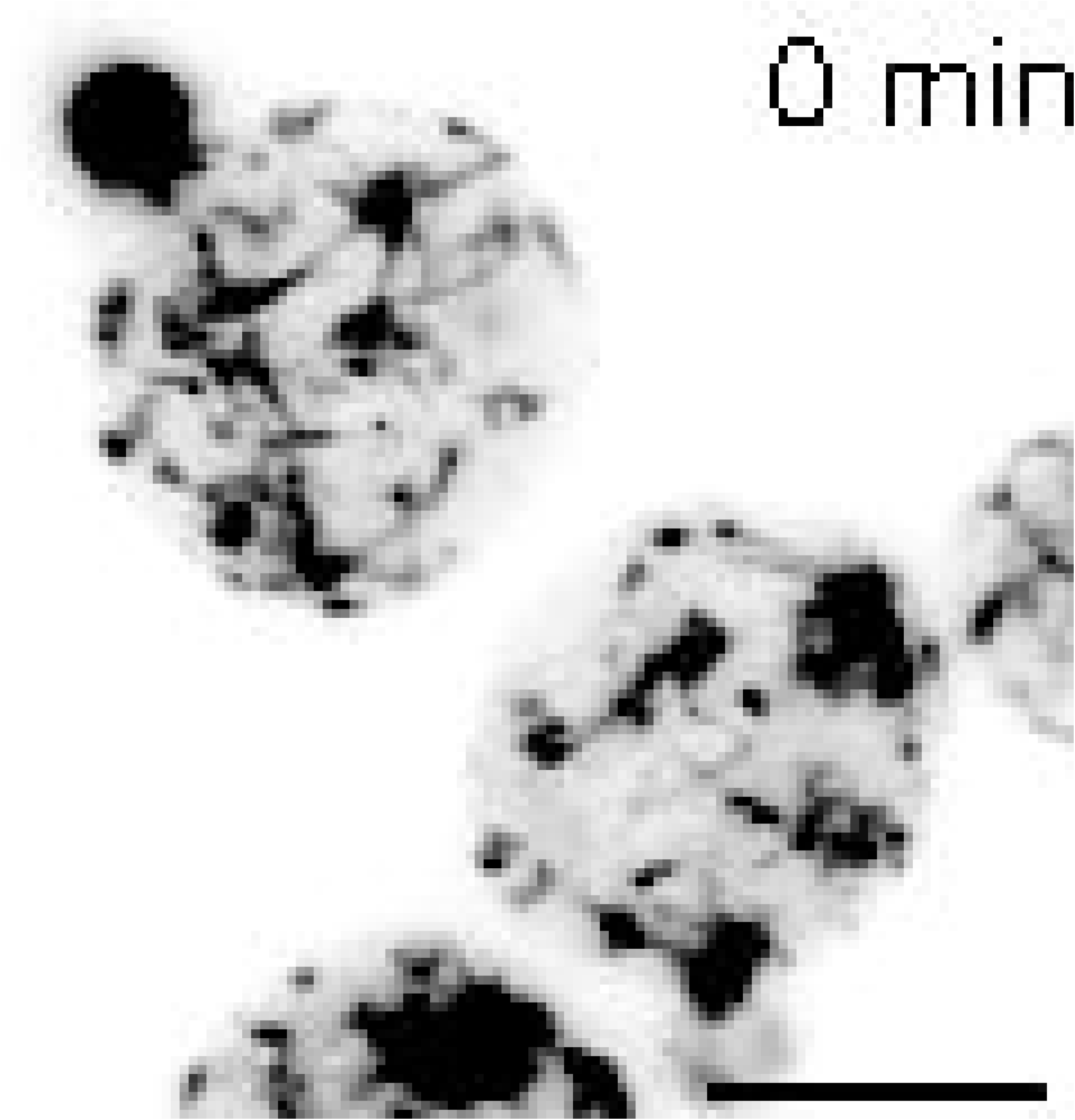

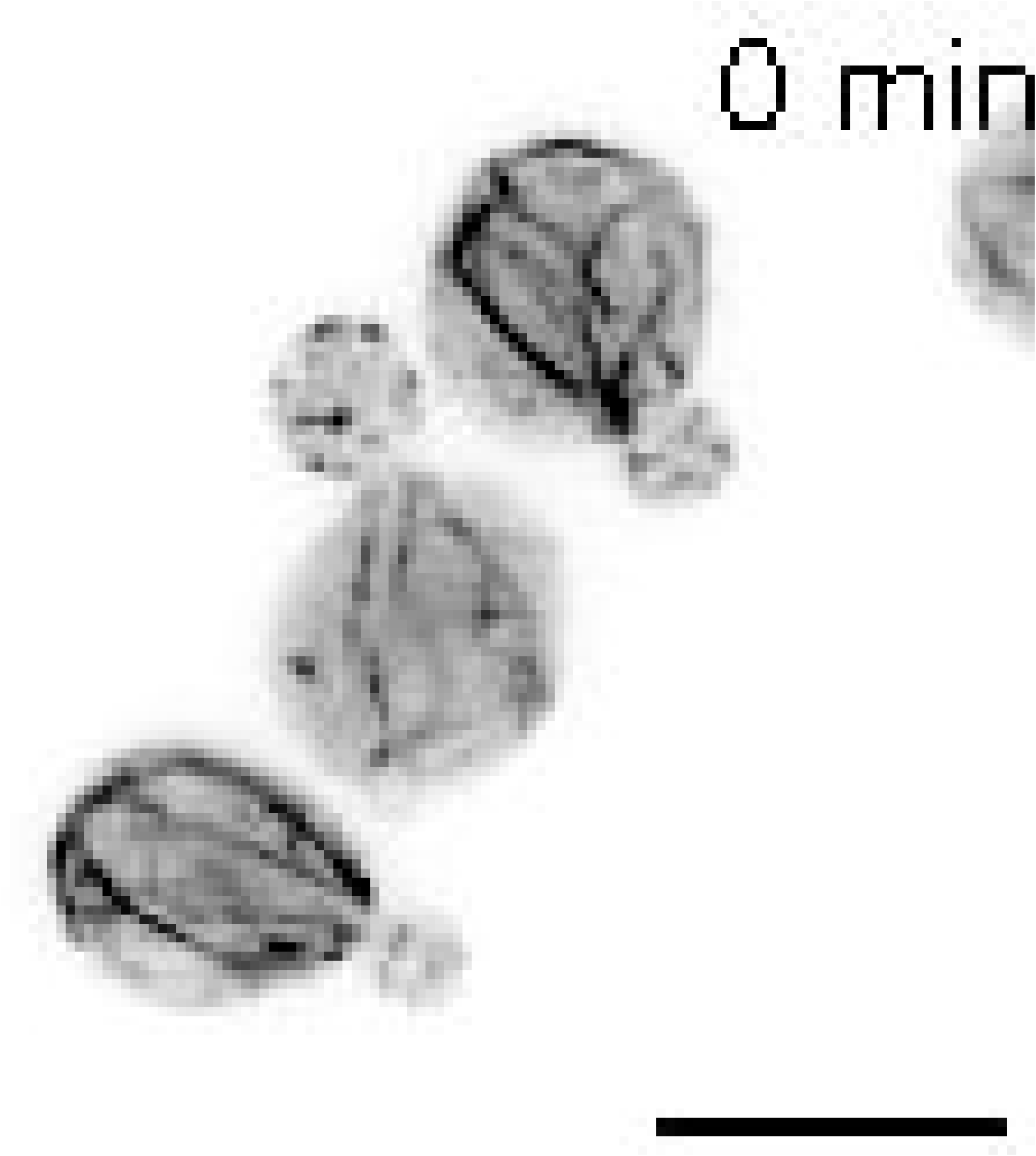

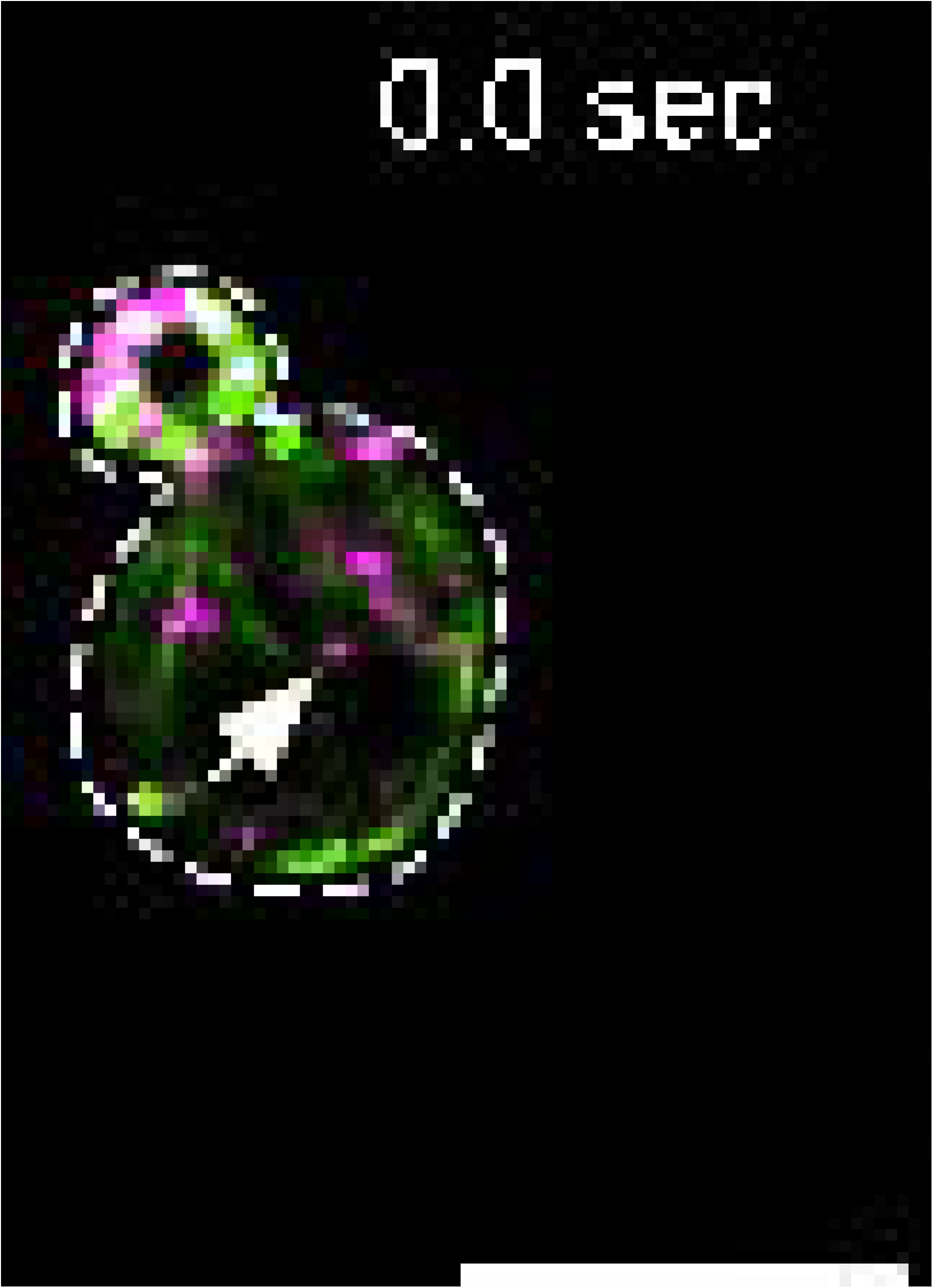

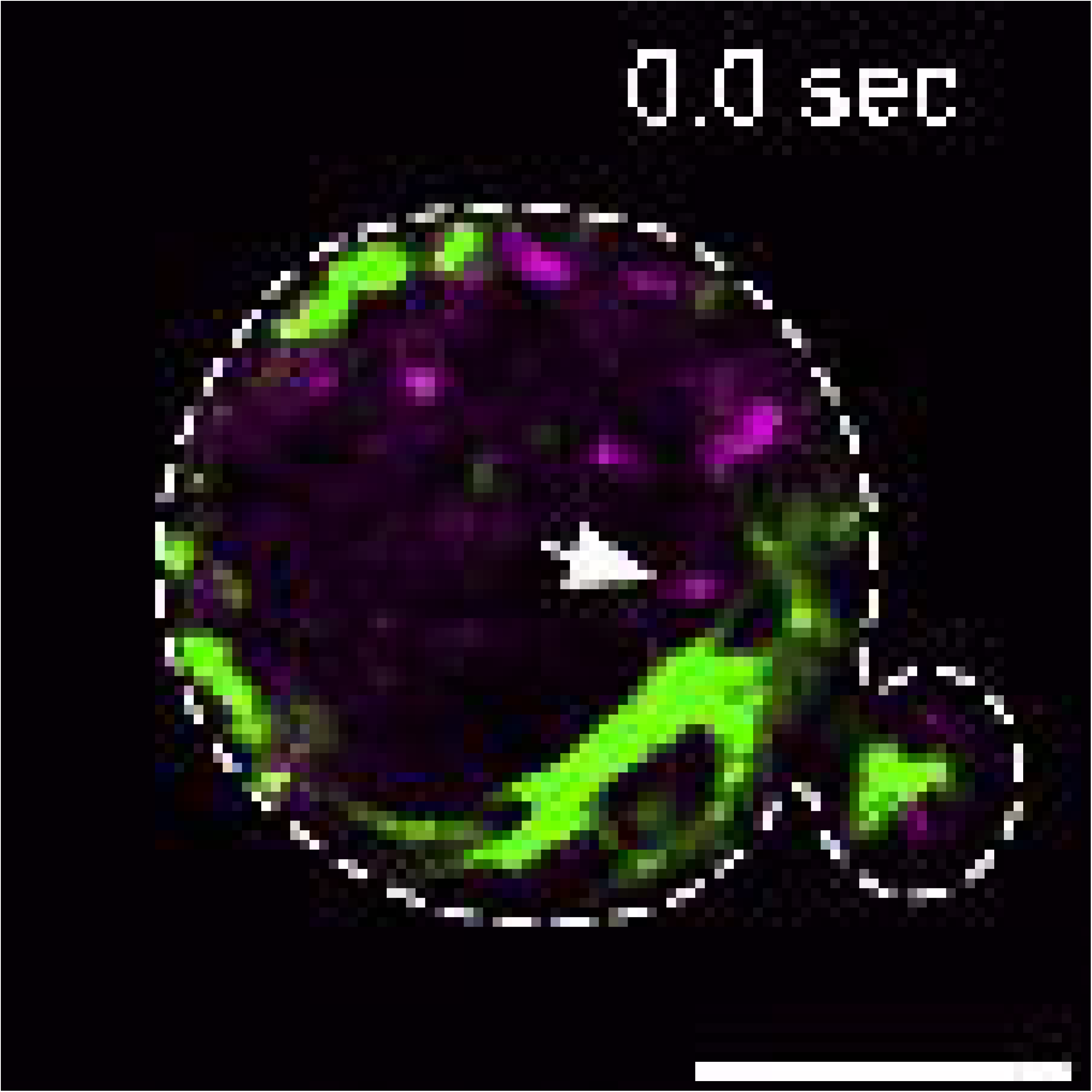

